# A unified approach for characterizing static/dynamic connectivity frequency profiles using filter banks

**DOI:** 10.1101/706838

**Authors:** Ashkan Faghiri, Armin Iraji, Eswar Damaraju, Jessica Turner, Vince D. Calhoun

## Abstract

Studying dynamic functional connectivity (dFC) has been the focus of many studies in recent years. The most commonly used estimator for dFC uses a sliding window in combination with a connectivity estimator such as Pearson correlation. Here, we propose a new approach to estimate connectivity while preserving its full frequency range and subsequently examine both static and dynamic connectivity in one unified approach. This approach which we call filter banked connectivity (FBC), implements frequency tiling directly in the connectivity domain contrary to other studies where frequency tiling is done in the activity domain. This leads to more accurate modeling, and a unified approach to capture connectivity ranging from static to highly dynamic, avoiding the need to pick a specific band as in a sliding window approach.

First, we demonstrated that our proposed approach, can estimate connectivity at frequencies that sliding window approach fails. Next we evaluated the ability of the approach to identify group differences by using the FBC approach to estimate dFNC in a resting fMRI data set including schizophrenia patients (SZ, n=151) and typical controls (TC, n=163). To summarize the results, we used k-means to cluster the FBC values into different clusters. Some states showed very weak low frequency strength and as such SWPC was not well suited to capture them. Additionally, we found that SZs tend to spend more time in states exhibiting higher frequencies and engaging the default mode network and its anticorrelations with other networks compared to TCs which spent more time in lower frequency states which primarily includes strong intercorrelations within the sensorimotor domains. In summary, the proposed approach offers a novel way to estimate connectivity while unifying static and dynamic connectivity analyses and can provide additional otherwise missed information about the frequency profile of connectivity patterns.

## Introduction

Functional connectivity and its cross-network analog, functional network connectivity (FNC) have been the focus of many neuroimaging studies over the past few decades. As the methods used to estimate FNC can be used to estimate FC and vice versa in most cases, in this article we will use the term FNC when talking about connectivity which includes FC too.

Methods developed to estimate FNC can be grouped into two major categories: those that assume connectivity among different networks of the brains is constant through time (static FNC; sFNC) and those that assume temporal variation in connectivity (dynamic FNC; dFNC). sFNC and dFNC approaches have proven to be extremely informative about both healthy (Allen, et al., 2014; Liegeois, et al., 2019; Vidaurre, et al., 2017) and disordered brain function (Damaraju, et al., 2014; de Lacy, et al., 2017; Jin, et al., 2017; Kaiser, et al., 2016).

### dFNC estimation approaches

While static connectivity has resulted in many interesting findings (Di Martino, et al., 2008; van den Heuvel and Hulshoff Pol, 2010), this view of connectivity is limited to the average connectivity patterns over the entire experiment. Approaches designed based on dFNC relax the assumption of static connectivity. The most common way to estimate dFNC uses a sliding window to estimate time-varying connectivity. Typically, a sliding window is paired with Pearson correlation (SWPC) to estimate time-varying connectivity (Allen, et al., 2014; Faghiri, et al., 2018; Hutchison, et al., 2013b; Kucyi and Davis, 2014). Using a sliding window to estimate dFNC has the benefit of being straightforward but has two major shortcomings. First, we need to choose a window size for any sliding window approach. We want to choose a window size that is large enough so that the standard error is as small as possible. At the same time, the window size should be small enough to allow us to detect faster changes in dFNC (Hutchison, et al., 2013a). The second shortcoming of the sliding window approach is its low-pass nature, which has been reported previously (Leonardi and Van De Ville, 2015; Sakoglu, et al., 2010; Thompson and Fransson, 2015). This tells us that, regardless of the chosen window size, the estimated dFNC is subjected to a low-pass filter and therefore the full frequency range of dFNC is not captured. This may be the reason that Shakil et al. found that using a constant window size for SWPC is not a reliable solution to study dynamic connectivity (Shakil, et al., 2018). Note that sliding window can also be used with other estimators such as multiplication of temporal derivatives (Shine, et al., 2015) and weighted average of shared trajectory (Faghiri, et al., 2020; Faghiri, et al., 2019). A sliding window will also act as a low pass filter if used with these and any other estimators. For a detailed comparison of various dFNC estimators see (Xie, et al., 2019).

Another category of methods that aim to explore the frequency profile of connectivity uses time-frequency analysis ideas. The most well-known methods in this category utilize wavelets (Mallat, 1999). While these methods have resulted in many interesting findings in functional magnetic resonance imaging (fMRI) (Chang and Glover, 2010; Yaesoubi, et al., 2015; Yaesoubi, et al., 2017), these studies have several limitations. Firstly, the interpretation of the results in these studies can be challenging. The Chang and Glover implementation of wavelet resulted in a large amount of information without a way to succinctly summarize the results (Chang and Glover, 2010). To remedy this, Yaesoubi et al. proposed an approach using wavelets that can be used to study group differences, but their results are all in the wavelet domain (Yaesoubi, et al., 2015). This presents a difficulty in comparing the results to other dFNC studies as most dFNC studies work in the time domain (the time domain results are typically considered easier to interpret). In addition, and perhaps more importantly, both wavelet approaches perform frequency tiling in the activity domain instead of the connectivity domain. We believe that to discuss the frequency properties of dFNC it is important to implement all time-frequency tiling steps directly in the connectivity domain. The reason behind this statement is that the relationship between the activity and connectivity domains is unknown (and possibly non-linear); therefore, the frequency information is distorted when transforming from the activity to the connectivity domain.

Apart from the two categories of approaches mentioned above, other methods have been proposed that do not estimate dFNC directly but rather explore different aspects of connectivity dynamics. These methods include hidden Markov models (Ou, et al., 2013), coactivation patterns using clustering approaches (Liu and Duyn, 2013), and window-less dictionary learning approaches (Yaesoubi, et al., 2018).

### Schizophrenia-related findings

There have been a number of studies of functional connectivity in schizophrenia individuals (SZ) using resting fMRI. For example, Camchong et.al. reported hyperconnectivity between default mode network (DMN) and the rest of the brain (Camchong, et al., 2011). In another work, Jafri et al. used a whole brain approach to study the differences between typical controls (TC) and SZ (Jafri, et al., 2008). They reported that SZ showed increase connectivity between DMN and visual and frontal functional domains compared to TC. Damaraju et al., reported that SZ compared to TC show increased connectivity between thalamus and sensory functional domains (Damaraju, et al., 2014). Damaraju et al., also reported decreased static connectivity in sensory domains when comparing SZ to TC. Decrease connectivity in SZ compared to TC has been reported in other studies too (Dong, et al., 2018; Erdeniz, et al., 2017; Friston and Frith, 1995; Lynall, et al., 2010; Skudlarski, et al., 2010).

A few studies have evaluated the dynamic aspect of connectivity in SZ population. Using SWPC, Damaraju et al., found that SZ compared to TC tend to stay less in states which shows strong overall connectivity while they tend to spend more time in states showing weak connectivity between different domains (Damaraju, et al., 2014). Other studies also reported transient reductions in both functional connectivity and networks activities (Iraji, et al., 2019a; Iraji, et al., 2019b). Miller et al., reported less dynamism (e.g. less change in transient connectivity patterns) in SZ compared to TC (Miller, et al., 2016). For a recent review on connectivity related findings (both static and dynamic findings) in SZ population please see (Mennigen, et al., 2019).

A different set of studies have explored the spectral properties of fMRI time series between SZ and TC. An earlier study found that the frequency profile of default mode is altered in SZs compared to TCs (Garrity, et al., 2007). In addition, Fryer et al. used voxel wise amplitude of low-frequency fluctuations (ALFF) between frequencies 0.01-0.08 to find that SZs have lower ALFF compared to TCs especially in posterior cortex, occipital and cerebellar lobes (Fryer, et al., 2016). This observation has been reported in other studies as well (Alonso-Solis, et al., 2017; Calhoun, et al., 2012; Hare, et al., 2017; Hoptman, et al., 2010). On the other hand, Alonso-Solis et al. report that ALFF for SZ is higher than that of TCs in insula (Alonso-Solis, et al., 2017). So it seems reasonable to think that the relationship between ALFF differences in SZs and TCs are different for different regions. In addition, Yu et al. found that these differences are dependent on the frequency and future studies should consider studying different frequency bands (Yu, et al., 2014).

All these studies point to differences in the frequency profile of activity level time series in individuals with SZ. A natural evolution of these studies is to explore the frequency profile of connectivity level information. An earlier attempt at this utilized wavelet coherence methods (Yaesoubi, et al., 2017); But as mentioned earlier, this method implemented frequency tiling in activity space, therefore the relationship between frequency and connectivity patterns is not direct.

### Frequency tiling in the connectivity domain

In this work, we emphasize the importance of differentiating the connectivity domain frequency profile from the activity domain frequency profile (more on this in discussion). As mentioned previously, some studies have implemented frequency tiling in the activity domain and use theses tiles to make inferences about the connectivity frequency profile (which we believe is inaccurate). Here we proposed a new method to estimate connectivity which aims to implement frequency tiling directly in the connectivity domain. As a central part of this approach, we use filter banks for frequency tiling in the connectivity domain (unlike wavelet methods where frequency tiling is implemented in the activity domain). We do this without making any assumption about the frequency profile of connectivity (unlike SWPC where only low-pass connectivity values are estimated). Using filter banks, we are able to estimate the entire connectivity frequency spectrum, removing the direct impact of window size on connectivity estimates. In addition, our proposed approach enables us to examine specific frequency bands of connectivity which includes both static connectivity (connectivity at zero frequency) and dynamic connectivity (connectivity at non-zero frequencies) in one unifying approach. Finally, the results are in the time domain allowing us to compare our results with other studies more easily (unlike wavelet approaches where the final results are in the wavelet domain). To explain the benefits of our proposed approach we designed a toy example. Furthermore, to showcase the utilization of the proposed approach, we implemented it on a dataset used to study sFNC and dFNC previously (Damaraju, et al., 2014; Yaesoubi, et al., 2017). The reason we chose to use a dataset that has been studied extensively is that then we can situate our results in an existing rich literature, enabling us to compare results from different approaches.

### Removing the direct impact of window selection on connectivity

Another advantage of our approach (filter banked connectivity; FBC) is that this method removes the impact that incorrect window selection has on the estimated connectivity values. The reason behind this statement is that here we are looking at the full spectrum of connectivity instead of limiting ourselves to any given band without strong prior knowledge. To drive this point home, we did a very simple simulation with two different situations. Assume we have 2 different pairs of time series each extracted from different locations in the brain. The first pair have low frequency connectivity while the second pair have high frequency connectivity. Figure 1 illustrates this simulation. For the first scenario, both SWPC and low-pass FBC have estimated the ground truth very nicely (both have high correlation values with the ground truth). But for the second scenario, only high pass FBC has a high correlation with the ground truth. Therefore, in this situation, the connectivity information is lost if we use SWPC.

**Figure 1.**
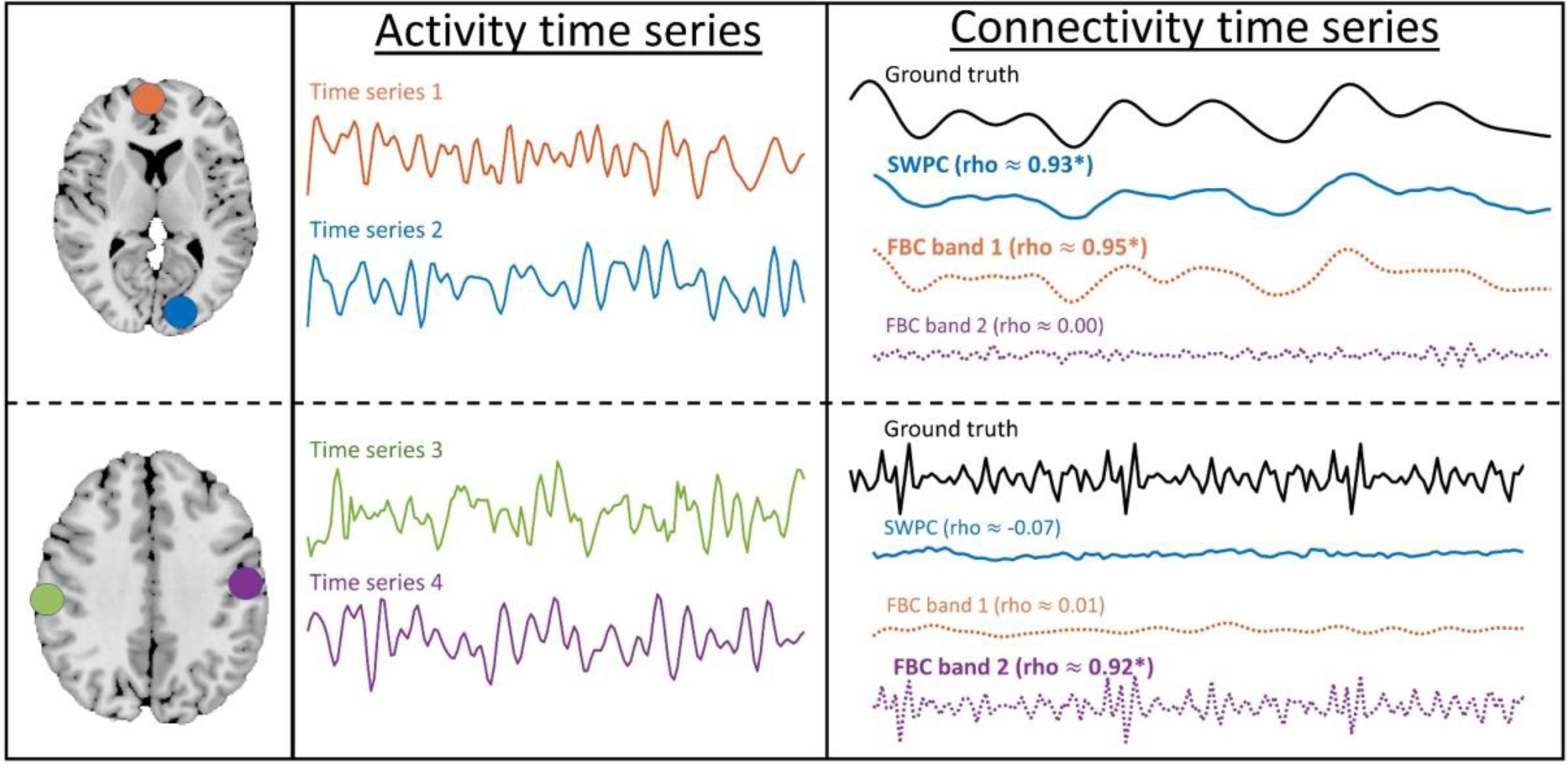
A toy example, to demonstrate the benefits of our approach. Each box demonstrates one specific situation. The left-most column, show the spatial maps of the simulated time series pair (this is just for demonstration). The middle columns show the activity times series pair themselves. The true and estimated connectivity for all methods is shown in the rightmost column. The correlation between each is estimated time series and the ground truth is brought in the parentheses here. In the top box, connectivity has a low frequency and both SWPC and FBC have managed to estimate it (FBC in its first band estimates). While in the bottom box, connectivity has a higher frequency. Therefore SWPC has not managed to estimate the connectivity while FBC has estimated it (in its second band this time).

In the methods section, we first introduce the proposed approach and its formulation. Next, we attempt to provide intuition into how our method performs using a toy example. After this the dataset used in the paper is introduced briefly. We mentioned findings in the results section and explored these in more detail in the discussion. Finally we discuss limitations and end the paper with some concluding remarks.

## Materials and Methods

### Filter banked connectivity

Assume we have two time series *x*(*t*) and *y*(*t*) where t is time. Centered SWPC at each time point (*r*_*x,y*_(*t*)) can be estimated as follows:

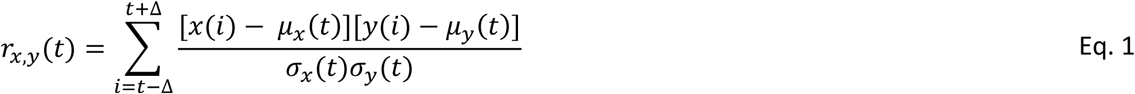

Where 2Δ + 1 is window size and *μ*_*x*_(*t*) and *σ*_*x*_(*t*) are windowed sample mean and windowed standard deviation respectively (for time series x) respectively. Their definitions are as follows:

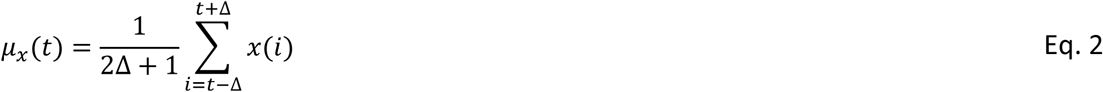

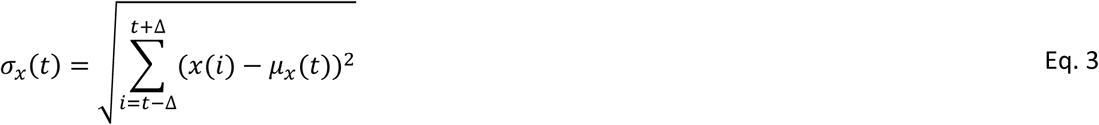

Similar equations can be used to estimate *μ*_*y*_(*t*) and *σ*_*y*_(*t*).

Now if we define two times series *h*(*t*) and *w*(*t*) such that:

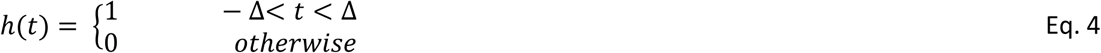

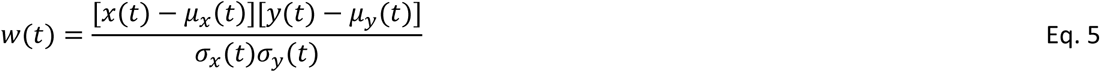

The convolution between these two time series can be written as:

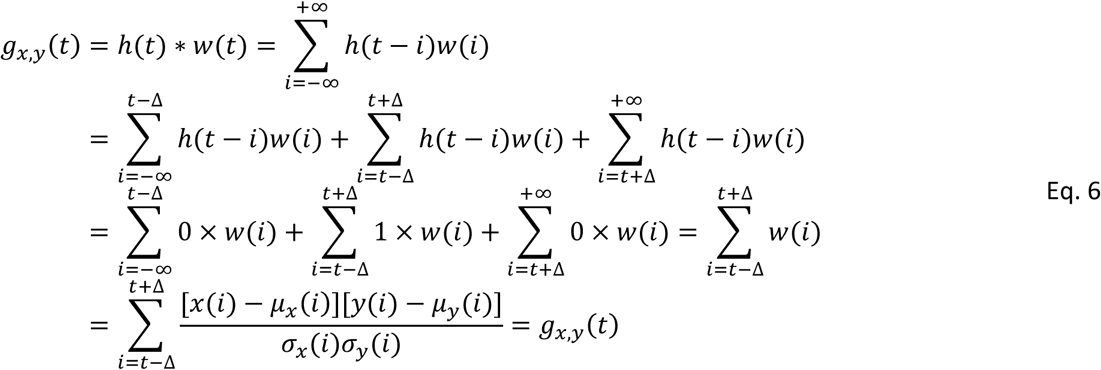

From the *g*_*x,y*_(*t*) equation we can see that it is quite similar to the equation for *r*_*x,y*_(*t*) in Eq. 1. Their difference is in how the windowed mean and standard deviation is calculated. In SWPC equation (Eq. 1) for each window we have one mean and standard deviation (the index of *μ*_*x*_ is t not *i*). In contrast, in *g*_*x,y*_(*t*) (Eq. 6) windowed mean and standard deviation are calculated using a window around each sample (the index of *μ*_*x*_ is *i* here). Therefore, we can interpret the convolution between *h*(*t*) and *w*(*t*) as an approximation for SWPC. Based on the system and signal theorem (Oppenheim, 1999), we know that the output of a system with an impulse response function *h*(*t*) and input of *w*(*t*) is *h*(*t*) ∗ *w*(*t*). So *g*_*x,y*_(*t*) (and SWPC that it approximate) is the output of a system with impulse response *h*(*t*) and input of *w*(*t*) (Figure 2.a). The defined *h*_*t*_ for SWPC system is a rectangular window that can be viewed as a low-pass filter. In other words, the output of the system (SWPC estimation) is a low-pass signal and we lose the high-frequency information of the connectivity.

**Figure 2.**
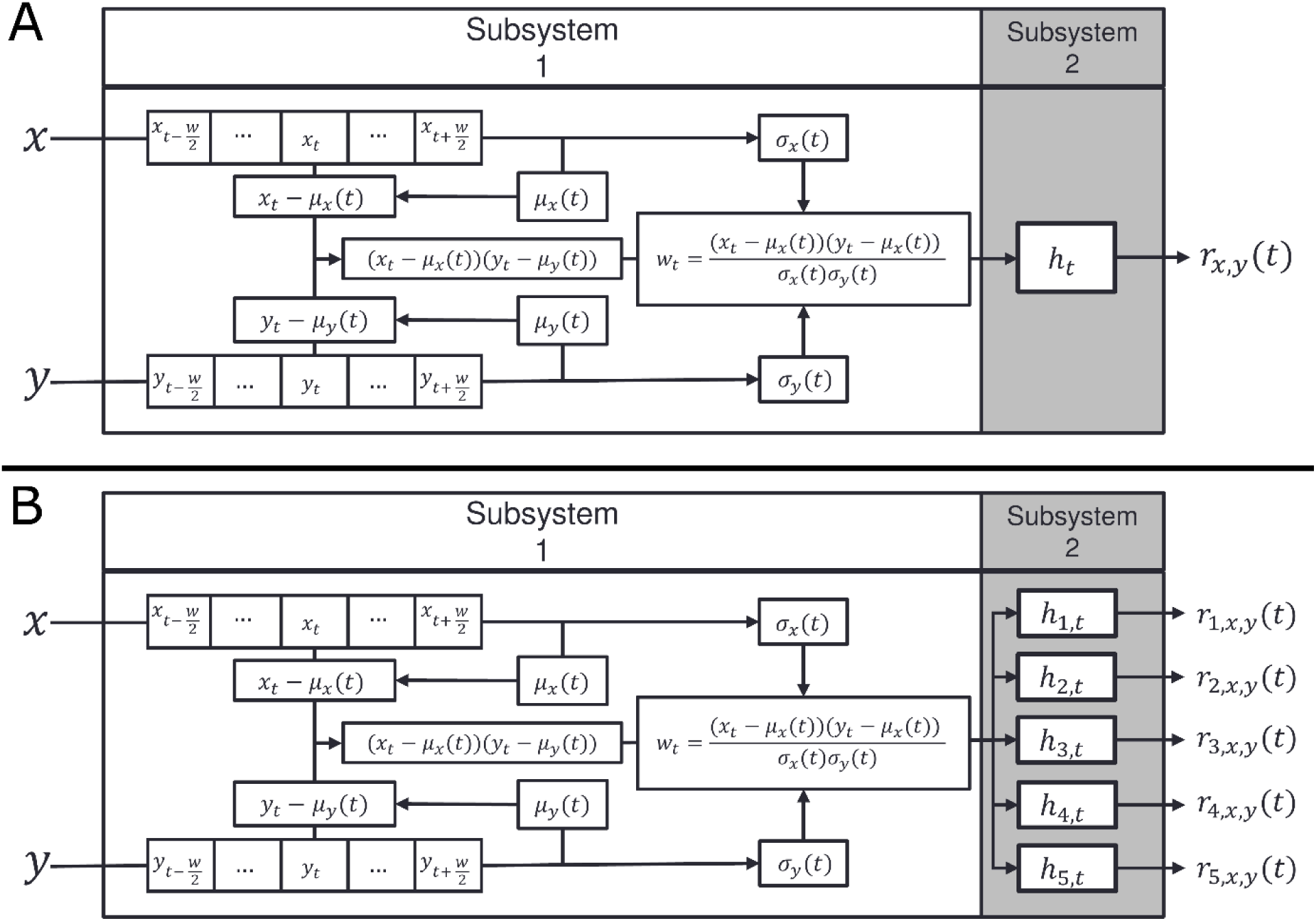
SWPC and FBC systems. A) SWPC system. B) FBC system. The subsystem 1 is shared between both SWPC and FBC. This subsystem uses a pair of time series and transforms activity space to connectivity space (*w*_*t*_ belongs to this space and has connectivity information). The difference between SWPC and FBC is in their subsystem 2. SWPC uses a low-pass filter to calculate a low pass version of *w*_*t*_ while in FBC instead of a low-pass filter, an array of filters are used. These filters, include (but are not limited) to the low-pass band examined in SWPC. FBC is more flexible in the sense that it does not make any assumption about the connectivity frequency and effectively spans a range of window sizes.

A filter bank is an approach that is used frequently in the electrical engineering field (Boashash, 2015). The basic idea behind a filter bank is to design an array of systems to filter one time series into its different frequency sub-bands (usually non-overlapping bands that cover the entire frequency spectrum).

In our proposed approach, we replace *h*(*t*) of SWPC (Figure 2.a) with a filter bank (Figure 2.b). Each filter in the designed filter bank has its own response function *h*_*n*_(*t*) where n represents filter index. In FBC instead of one low-passed connectivity time series (i.e. *r*_*x,y*_(*t*) as in SWPC), we have N time series each an estimate of connectivity in the sub frequency bands defined by *h*_*n*_(*t*). In other words:

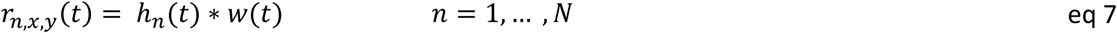

In this paper, we used Chebyshev type 2 filters. These filters are Infinite impulse response filters that have better frequency features compared to finite impulse response filters (Rabiner, et al., 1974). The issue with Infinite impulse response filtering is that it has nonlinear phase (compared to linear phase for finite impulse response filters), but because of the offline nature of fMRI data analysis we can use forward-backward filtering to achieve zero-phase filtering in all our analysis (Mitra and Kuo, 2006; Oppenheim, 1999). To get the optimal order for filters we used cheb2ord as implemented in MATLAB software to achieve at most 30 db attenuation in stopband and 3 db in the passband (Rabiner and Gold, 1975).

### Toy example

We have designed a simple toy example to provide insight into how our method works. Note that the aim for designing this toy example was not to comprehensively evaluate FBC for use with fMRI data but rather to convey intuition. In addition, we believe this approach can be used to analyze many types of datasets and therefore we designed the toy example to be a general illustration.

For this toy example, we use a multivariate normal distribution to simulate 6 time series:

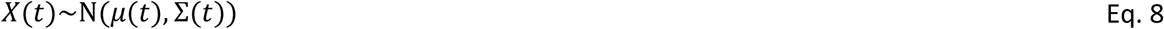

For simplicity, we put *μ*_*t*_ equal to 0 and each time series variance equal to 1. Therefore, the covariance matrix can be written as:

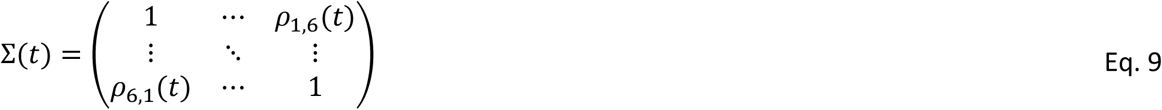

Where *ρ*_*i,j*_(*t*) is the correlation (i.e. connectivity) between time series *i* and *j* at time *t. ρ*_*i,j*_(*t*) has the form *ρ*_*i,j*_(*t*) = *A*_*i,j*_(t)cos (2*πf*_*corr*_*t*) where *A*_*i,j*_(t) forms the 6×6 matrix *A*(*t*). For all toy examples, we simulated time series using two connectivity states each with a length of 10,000 time points. Each state has a unique *A*(*t*) and *f*_*corr*_ where *f*_*corr*_ determines the frequency of connectivity while A determines its amplitude. Figure S1 shows the *A*(*t*) of the two states used in all scenarios. The time series starts with state 1 and switch to state 2 after 10,000 timepoints. The total length of the simulated time series is 20,000.

Two scenarios with different values for *f*_*corr*_ and analysis parameters were designed. In both scenarios, we use two filter banks (one low-pass filter and one band-pass filter). In the first scenario, we choose SWPC window size such that it covers the same frequency band as the 2 designed filters. This scenario represents a case where the window size is chosen correctly for SWPC (it covers all the connectivity frequencies where the information resides). In the second scenario, the SWPC only covers the low pass filter in the filter bank. This scenario represents a case where the window size is chosen larger than what should be used (i.e. we are filtering out some of the relevant connectivity frequencies). This scenario can happen either because of the researcher’s mistake (note that we generally do not know the ground truth about real data), or the technical limitation of the connectivity estimator paired with sliding window. For example, if sample Pearson correlation is used with sliding window (as is the case here), we know that if the number of samples is below a specific number, sample Pearson correlation estimators fails (it gives very skewed results and in the worst case estimates only 1 and −1). Therefore, there is a lower bound on window size if we use this estimator. As the frequency of a rectangular window is tied directly to its window size, the lower bound on window size causes a higher bound on frequency, i.e. the connectivity is low-passed.

To compare FBC with SWPC method, we analyzed the toy example data using both methods. To provide a more direct comparison between FBC and SWPC, the filters designed for toy examples did not cover the whole spectrum of the simulation (we cannot have a window that filters the whole frequency and reach a good estimation of correlation for SWPC). In addition, for simplicity purposes, we used the same window size for both FBC (window used to estimate sample mean and standard deviation) and SWPC. The results of both FBC and SWPC were then clustered into 4 clusters using k-means approach. The reason for choosing 4 as the number of clusters was because we had 2 original states. Each state has two extreme correlation value (+*A*_*t*_ and −*A*_*t*_ because of the sinusoid nature of *ρ*_*i,j*_) so we essentially have two pairs of states.

The *f*_*corr*_ value for each state and the designed filters frequency response will be shown when discussing the results.

### Real dataset and preprocessing

To demonstrate the utilization of FBC, we used it to analyses a resting state fMRI dataset including SZ and TC individuals. The data was obtained as a part of the Functional Imaging Biomedical Informatics Research Network (fBIRN) project (Potkin and Ford, 2009). The dataset used in this paper includes 163 TC and 151 SZ. The data acquisition and preprocessing steps are explained in our previous work (Damaraju, et al., 2014). To summarize, echo planar imaging was used to acquire 162 volumes of bold data at seven sites all using 3T MRI scanners. All scans were acquired using 2 sec as TR. Subjects’ eyes were closed during the scanning session.

Preprocessing was started with motion correction, slice-timing correction, and despiking. Next, data were registered to a common Montreal Neurological institute (MNI) template and smoothed to 6mm full width at half-maximum. For the last step of preprocessing, each voxel time series was variance normalized.

To decompose the data into 100 spatially independent time series and their associated spatial maps the pipeline proposed by Allen et al., was used (Allen, et al., 2014). In the proposed approach group spatial independent component analysis (GICA) implemented in the GIFT (http://trendscenter.org/software/gift) software (Calhoun, et al., 2001b; Erhardt, et al., 2011a). The 162 time points for each subject were first reduced into 120 dimensions using principal component analysis (PCA). All subjects’ reduced data were then concatenated and another PCA was used to reduce the dimension to 100. Finally, independent components were estimated using infomax algorithm (Bell and Sejnowski, 1995). ICA was repeated 20 times in ICASSO algorithm (Himberg and Hyvarinen, 2003) and the most central solution was selected for stability purposes (Du, et al., 2014). Subject specific time series and their associated spatial maps were calculated using a back reconstruction approach (Calhoun, et al., 2001a; Erhardt, et al., 2011b). The spatial maps of these 100 components were visually inspected and 47 components were chosen as components of interest and were grouped into 7 functional domains. These time series where then band-pass filtered between 0.01-0.15 Hz using Butterworth filter (5th order). The data used for the current project are the same 47 components used by Damaraju et al. (Damaraju, et al., 2014). The 7 functional domains are: auditory (AUD), attention/cognitive control (CC), sub-cortical (SC), Cerebellar (CB), default mode (DM), sensorimotor (SM) and visual (VIS). The spatial maps of all the components included in each functional domain can be viewed in Figure S2.

### FBC and SWPC analysis (real data)

To analyze the fBIRN dataset using FBC pipeline, each component pair (from the pool of 47 components) was used to calculate a specific *w*_*t*_ (Eq. **5**) for that pair. For calculating *w*_*t*_, a window with size equal to 10TR (22 s) was used. This step resulted in 1,081 *w*_*t*_ time series (47 × (47 − 1)/2). All *w*_*t*_ were then passed through our designed filter banks. For this paper, 10 IIR filters were used to filter all the *w*_*t*_ values. To make sure the phase of *w*_*t*_ is not affected by the filters, we used a forward-backward filtering approach (Mitra and Kuo, 2006). The filter bands frequencies are:

Band 1: 0.000 - 0.025 Hz

Band 2: 0.025 – 0.050 Hz

Band 3: 0.050 - 0.075 Hz

Band 4: 0.075 - 0.100 Hz

Band 5: 0.100 - 0.125 Hz

Band 6: 0.125 - 0.150 Hz

Band 7: 0.150 - 0.175 Hz

Band 8: 0.175 - 0.200 Hz

Band 9: 0.200 - 0.225 Hz

Band 10: 0.225 - 0.250 Hz

The filtered values resulted from all filter banks were then clustered using k-means approach. For k-means clustering, we used K-means++ algorithm (Arthur and Vassilvitskii, 2006) with Squared Euclidean distance. The clustering was repeated 30 times with different initial cluster centroids and the one with the lowest within cluster distance was selected as the best results.

Next, we calculated how many time points each subject has spent in each specific state (fraction rate). This value can be calculated for all 10 frequency sub-bands separately or for all of them combined. In summary, for each state, we produce a plot that is quite similar conceptually to the frequency response of the corresponding state. In addition, we have compared the fraction rate between TC and SZ for each state. Figure 3 illustrates the whole pipeline used including gICA and clustering steps.

**Figure 3.**
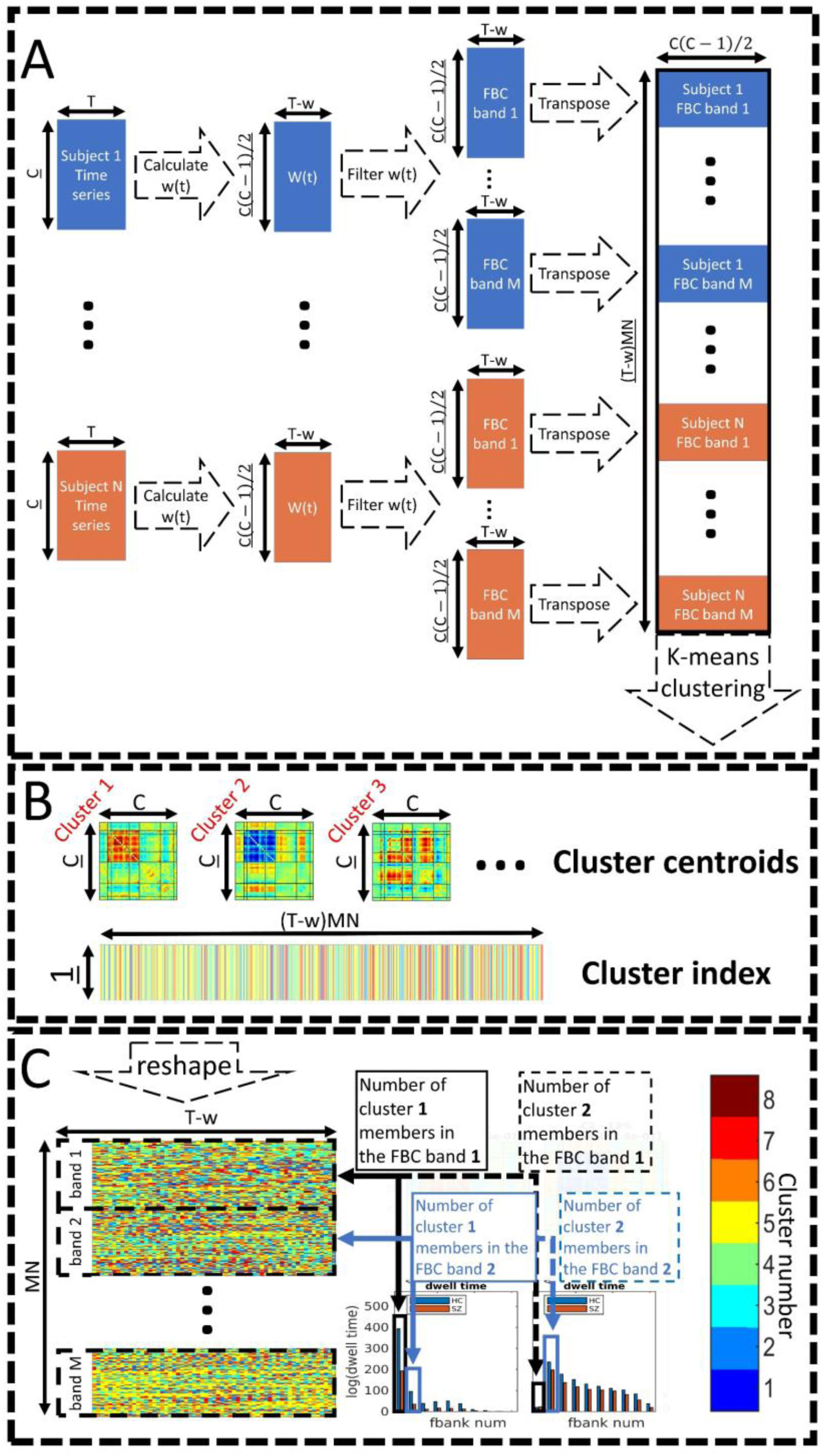
The whole pipeline used in this manuscript. A) FBC estimation. First, all subjects 47 time series were used to estimate w(t), resulting in 1081 time series for each subject. Then w(t)s were filtered into 10 bands and concatenated into one big matrix. K-means clustering was applied to this matrix (feature size 1081) which resulted in 8 clusters. B) the kmeans clustering resulted in 8 clusters. The cluster centroids were matrices with size 47 by 47. While the cluster index is a vector with time by subject_number by 10 (number of bands). C) This cluster index vector was reshaped into 10 matrices each with a size equal to time by subject number. Each of these 10 matrices belongs to one band and can be used to calculate the fraction rate for each band/ each cluster.

Within this pipeline, there are two analysis parameters that must be selected. First, we have to choose a window size which will be used to calculate *w*_*t*_ (i.e. calculate mean and std of component pairs). Here we have chosen a window size of 11 time points (22s) for this step. Results from other window sizes are provided in supplementary material, Figures S4 through S12. As seen in these figures, the results are similar across all window sizes. Secondly, the number of clusters (i.e. k) needs to be selected for k-means. We chose k = 8 as the desired number of clusters in all our analyses explained in this manuscript. Our selection was based on a calculated metric (for more details refer to the supplementary material; Figure S3). In addition, we also performed our analysis using different cluster numbers and provide this information in the supplementary material (Figures S24 through S12). All statistical tests were corrected for multiple comparison we used a method that controls false discovery rate (fdr) on the results (Benjamini and Hochberg, 1995).

## Results

As mentioned in the methods section, we designed several toy example scenarios using a multivariate Gaussian pdf to demonstrate the benefits of FBC compared to SWPC. In addition, we show the use of FBC on real data including SZ and TC.

### Toy example Results

All the toy example results are summarized in Figure 4. Each sub figure (boxes a through f) has 7 rows. Row numbers are brought in the left-hand side of boxes a and d. The first and fifth rows demonstrate the normalized frequency response of the covariance matrix for FBC (first row) and SWPC (fifth row). The second and third rows demonstrate the mean and standard deviation of the centroids calculated from the toy example data while the sixth and seventh rows demonstrate the same measures for SPWC. Conceptually speaking, the mean shows the estimated values are close to the true values on average while std represents the standard error of the estimators. The fourth row shows the total fraction rate of each state in relation to the two filters. This information is exclusive to FBC and is not available for SWPC (SWPC is essentially one filter). Cluster 1 and 2 are the maximum and minimum values for state 1 (connectivity in state 1 oscillate between these values). Cluster 3 and 4 are the maximum and minimum values for state 2. True amplitude matrices, *A*(*t*), for the two states are shown in Figure S1.

**Figure 4.**
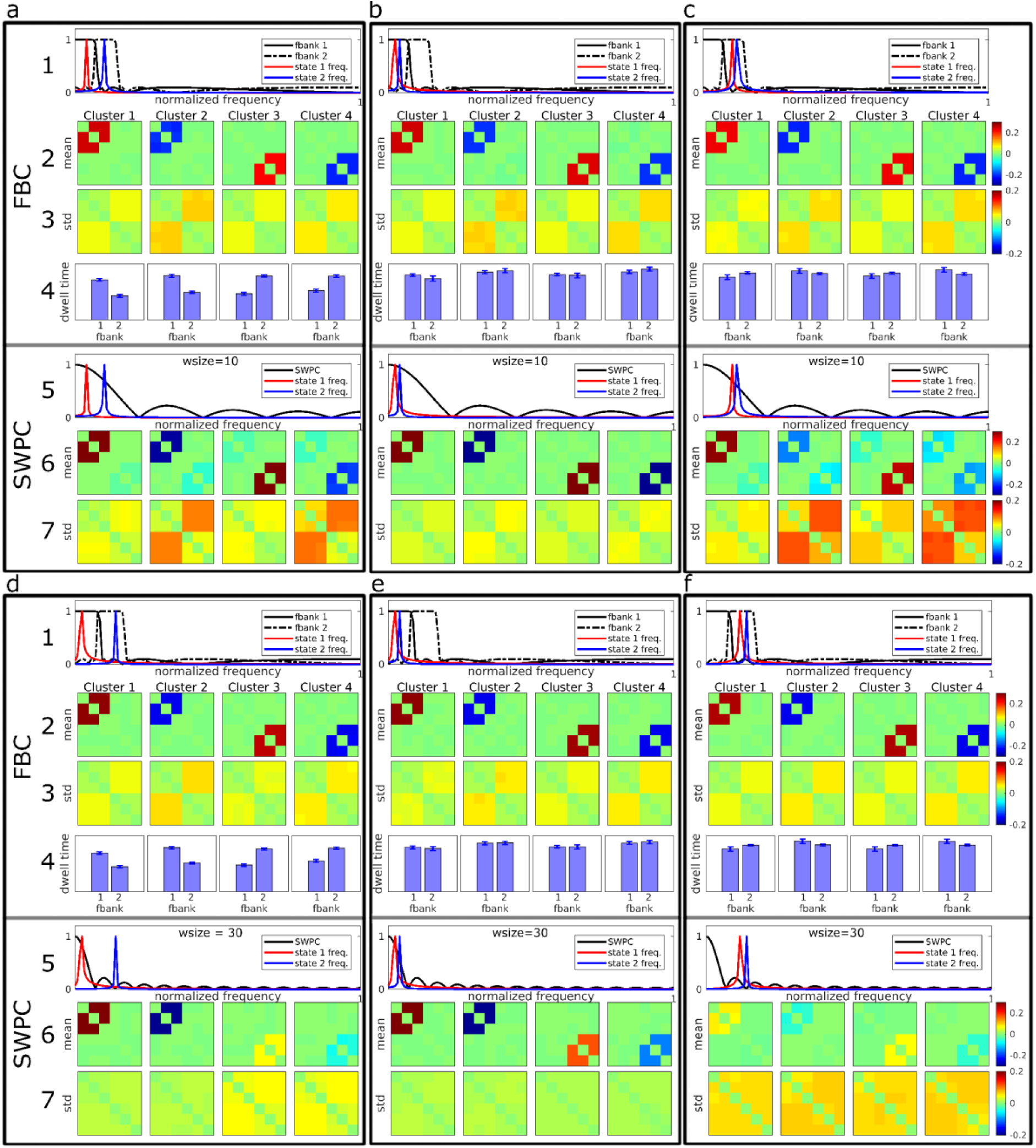
toy example results. Each figure (a through f) show the results from both FBC and SWPC analysis. The numbers show the row number for each box. Rows 1 through 4 illustrate FBC related results while the last three rows illustrate SWPC results. Rows 1 and 5 show the connectivity frequency of each specific scenario in addition to the frequency response of filter banks (1^st^ row) and the sliding window (5^th^ row) used for the analysis. Rows 2 and 6 show the mean of estimated cluster centroids (i.e. connectivity states). Rows 3 and 7 show the std of the estimated cluster centroids. The fourth row shows the frequency profile of the estimated cluster centroids for FBC (this information is exclusive to the FBC approach and is one of this method strengths). Figures a through c is the scenario where the SWPC sliding window is chosen correctly (connectivity frequencies are included in the main lobe of the sliding window for all three cases). In these cases, SWPC has managed to estimate states correctly in the three cases but when at least one of the states has higher frequencies where SWPC main lobe has a lower value (figure a and c), SWPC is not able to distinguish between different states well (the mean matrices in row 4 show both state patterns). In contrast, FBC has managed to estimate the 2 states very distinctly (e.g. the state 2 pattern does not appear in both clusters 1 and 2 in figure c unlike SWPC results). In addition, FBC is showing superior std (i.e. lower) in these cases. Figures d through f illustrate the results from the second scenario where SWPC window size is not chosen correctly (either because of investigator mistake or Pearson correlation technical limitations). Apart from the case where both state connectivity frequency is in the passband of SWPC (figure e), SWPC is not able to estimate the two states well (i.e. low values for means in row 6). In contrast, FBC does well in all three cases (high mean values). Apart from case e, where connectivity frequencies are very low and SWPC has an advantage over FBC, in the other 2 cases, std of FBC is superior too. One final note is that in the cases where the connectivity frequencies are in two separate bands (figures a and d) the connectivity frequency profiles resulted from FBC show that clusters 1 and 2 have lower frequency while clusters 3 and 4 have higher frequencies.

In the first scenario (Figure 4 a through c), the SWPC window size (10 time points) covers the same frequency band as the two filters used in FBC. This scenario includes three specific situations. In the first situation, each state *f*_*corr*_ is located in one separate filter (Figure 4a) while in the second and third situations *f*_*corr*_s are either in the first filter (Figure 4b) or in the second filter (figure 4c). Looking at the top three rows of Figure 4, we can make several observations. The means of estimated clusters are estimated very distinctly using FBC. i.e., cluster 1 and 2 only show state 1 *A*_*t*_ while cluster 3 and 4 only shows state 2 *A*_*t*_. In contrast, if we look at SWPC results, mean of estimated clusters are only estimated distinctly when both states have low frequencies (Figure 4b). When the frequencies of the states are higher (Figure 4a and c) cluster means are not distinct. For example, cluster means of Figure 4c for SWPC shows both state 1 and 2 *A*_*t*_ patterns (Figure S1) in all clusters.

In the second scenario, the window size is longer compared to the first scenario (30 time point) and only covers the frequency band of the first filter of FBC (Figure 4 d through f). This scenario represents the case where SWPC window size is not chosen correctly. i.e., the passband frequency of the SWPC window does not cover all frequencies where state connectivity frequencies are located. This can happen either because of user incorrect choice or because of the technical limitation of the estimator that is paired with a sliding window (Pearson correlation here). This is shown in Figures 4d and 3f where at least one of the states has connectivity frequency outside the passband of the SWPC window. Looking at SWPC results in Figure 4d we can see that cluster 3 and 4 show very weak versions of state 2 *A*_*t*_. In a more severe case where all states’ connectivity frequencies are outside the passband of SWPC (Figure 4f), all SWPC cluster mean values are very weak. Contrary to SWPC results, FBC mean values for cluster centroids are quite strong and similar to state 1 and 2 *A*_*t*_. Note that the scales of the images are the same across all scenarios.

In addition to the means, we can also interpret the std values too. Based on Figures 4 a through c we can see that SWPC produces higher std values compared to FBC std values for null connectivity elements (matrix entries where true connectivity is zero in each state). In contrast, for the second scenario results, we can see that using longer window sizes results in lower std values where the true connectivity frequency is very low. This is the case for Figure 4d (first state) and Figure 4e (both states). In addition, when the specific state has a higher frequency than the SWPC frequency band, the std values are noticeably higher for all matrices’ entries. This can be seen for the second state in Figure 4d and both state results in Figure 4f.

Another interesting observation here can be made for the case where the two states *f*_*corr*_s are located in separate filter bank frequency bands (Figure 4a and 3d). The fraction rate for FBC (row 4 in each box 3a and 3d) shows that cluster 1 and 2 spend more time in the first filter bank while the cluster 3 and 4 spend more time in the second filter. This is an accurate reflection of the ground truth and shows that FBC can provide correct frequency specificity in the case where connectivity frequencies are distributed in different filter frequency bands. This is also an exclusive and important feature of the FBC approach. This point will be expanded upon more in the discussion section.

Figure 5 shows the correlation between the estimated clusters and the true centroids (both positive and negative matrices shown in figure S1). Based on this figure we can see that in most cases FBC performs better (has a higher correlation) except the case where the connectivity frequencies are well inside the SWPC frequency window (cases b and e). Thus, if even one of the states has a higher frequency than what is covered by the SWPC window, FBC performs better (cases a, c, d, and f). In addition, compared to SWPC, FBC shows more robust performance; i.e. FBC performance is similar across different scenarios. This means that FBC performance is not impacted greatly by the true connectivity frequency which is unknown to us. Finally, FBC results seem to have less variation (spread of correlation values) compared to SWPC results which have only low variation in cases b and d. Note that these two cases are essentially the best-case scenarios for SWPC.

**Figure 5.**
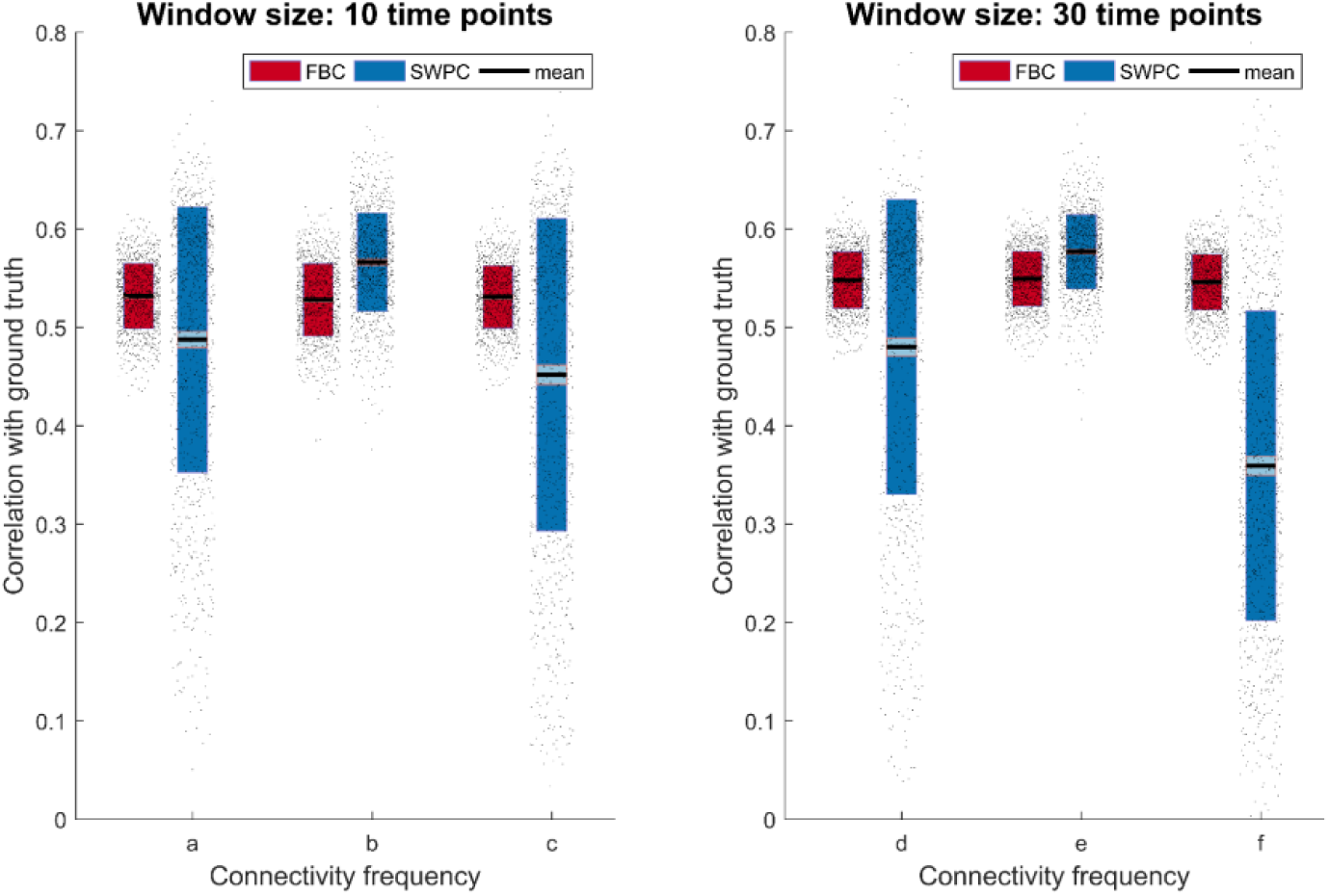
Correlation of estimated cluster centroids with ground truth for all toy examples. The left figures show the results for the first scenario (a through c; window size equal 10 time point) while the figure is for the second scenario (d through f; window size equal 30 time point). As can be seen here FBC performs better (i.e. have higher correlation) for most of the cases. Only in cases b and e, SWPC performs better than FBC (connectivity frequency is well within the band-pass of SWPC). In the cases where even one of the connectivity frequencies spread outside the SWPC window size (cases a, c, d, and f) FBC performs better than SWPC. Note also that the FBC mean Correlation values remain mostly the same for all 6 cases. This observation combined with the fact that we never know the correct window size, suggests FBC as a more robust solution.

### Group differences between TC and SZ in the fBIRN data

As mentioned in the method section, FBC was utilized to analyze the fBIRN data set. After calculating FBC values for all component pairs, we compute 8 clusters using the k-means method. The results can be seen in Figure 6. Based on the fraction rate values across all 10 bands (second row in each separate box) we can group the clusters into 3 groups: low-pass, band-pass and high-pass clusters.

**Figure 6.**
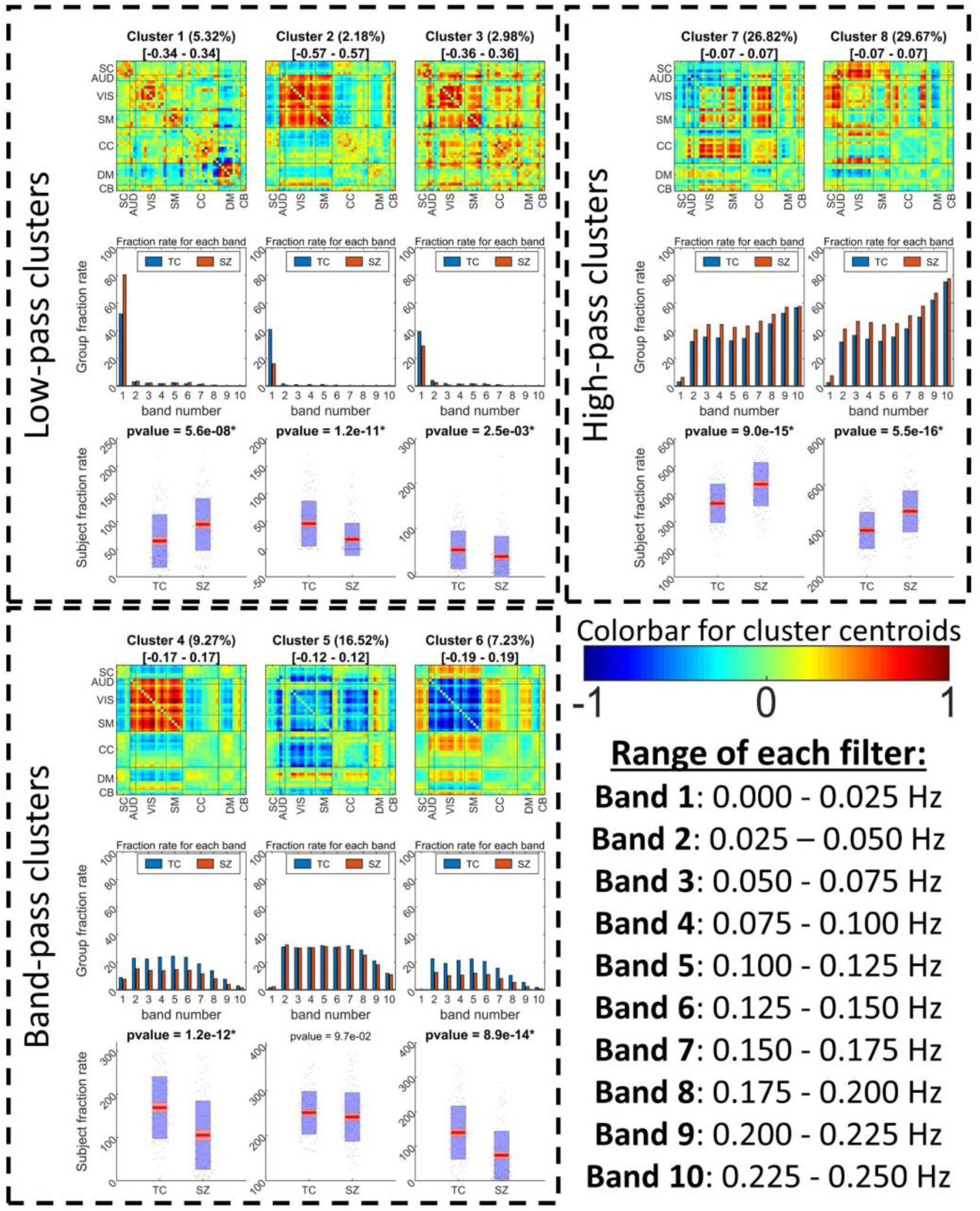
fBIRN FBC results. In each box, the first row shows the cluster centroids where the title gives the ratio of each state occurrence and range of each state centroid colormap (e.g. for C1 −0.34 is blue while 0.34 is red). The second row shows each cluster frequency profile (fraction rate of each band) for TC and SZ separately. The third row compares the fraction rate across all bands between TC and SZ (the title of these contains the comparison p-value where the significant ones are in bold font). The first observation we can make is that FBC has resulted in states which show opposite patterns to some other states. These opposite patterns are not visible within the SWPC results as some of them show a more high-pass frequency profile (possibly the reason these are not visible in the SWPC results). These opposite patterns might point to the presence of connectivity oscillation as designed in our toy examples. Another observation we can make is that states 2 and 3 show a very strong low-pass frequency profile where TCs tend to spend more in state 1 while SZ spend more time in state Interestingly, states 1 and 2 are are very similar to the static connectivity calculated from SZ and TC respectively (see Figure 7). States 7 and 8 is a finding exclusive to FBC which SZs tend to stay more in significantly more compared to TCs. These states show a very sparse connection in AUD/VIS/SM. This is in contrast with states 2 and 4 (higher fraction rate for TC) very strong connections in these domains.

Figure 7 shows all the SWPC results. Figure 7a depicts the static FNC (FNC calculated over the entire time series using Pearson correlation) for the TC and SZ groups separately. Looking at Figure 7a, we can see that the TC group has a stronger positive connectivity block in AUD, VIS and SM functional domains compared to SZ static FNC. Comparing these results to the 2 low-pass clusters in Figure 6 (Cluster 1 and 2), we can see a resemblance between static FNCs and Clusters 1 and 3. This can also be verified using Figure 7c where correlation is used to assess the similarity of the cluster centroids. Based on this figure, TC static FNC highest correlation is with FBC cluster 2 while SZ static FNC has a higher correlation with FBC cluster 1. Another observation that supports this comparison is that TCs spend significantly more time in Cluster 2 compared to SZs (Figure 6 last row in each box). In addition, SZs tend to stay more in FBC cluster 1 compared to TC.

**Figure 7.**
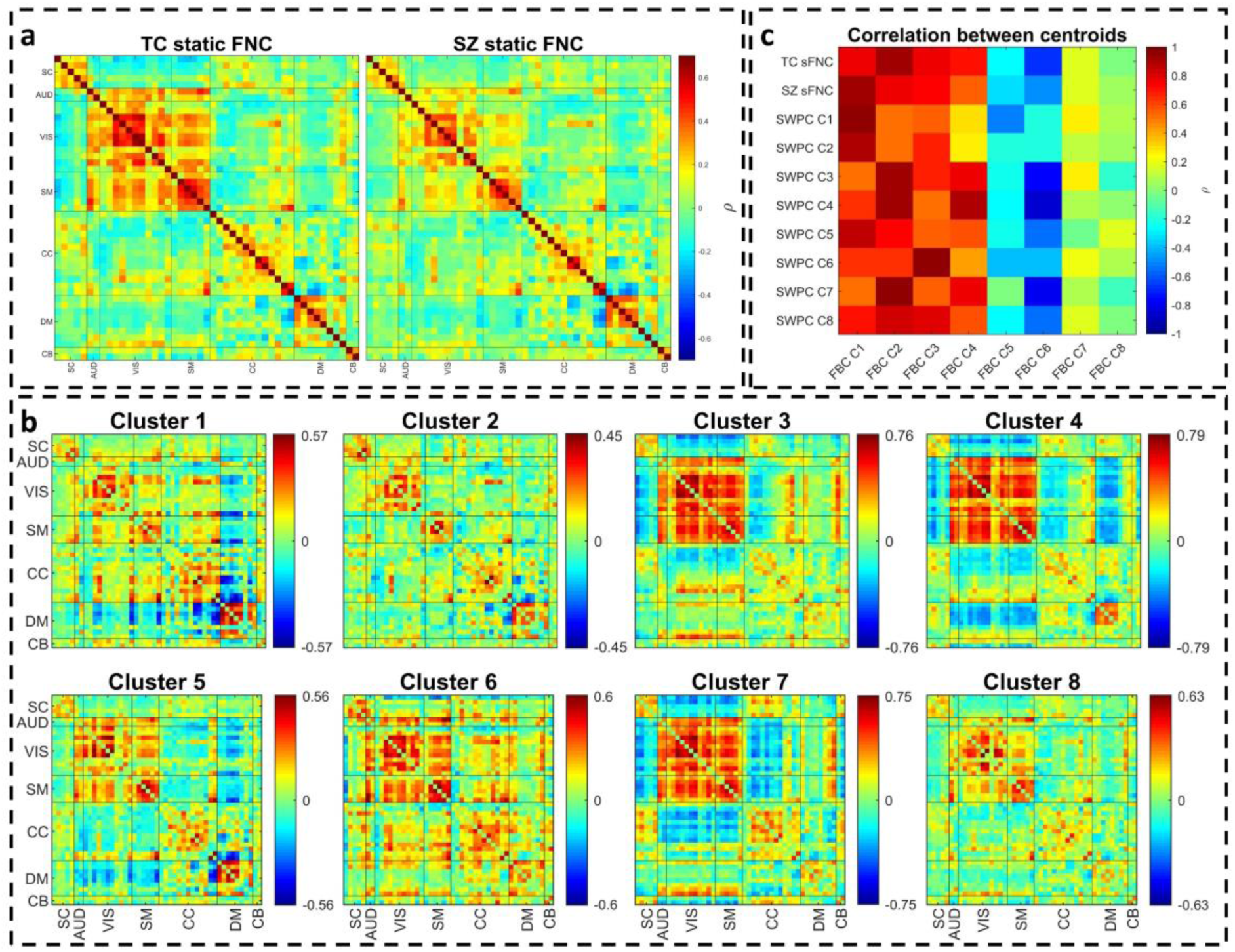
Pearson correlation results. a) static connectivity estimated using all the time points of the time series. b) estimating dFNC using SWPC and then clustering all the results into 8 clusters. c) correlation between Pearson correlation (both static and dynamic states) and FBC results. As can be seen In this figure, sFNC for SZ and TC are quite similar to FBC clusters 1 and 2 respectively (in c, they have the highest correlation). In addition, all SWPC clusters are quite similar to the first 4 clusters of FBC. FBC clusters 1, 2 and 3 are all low pass and cluster 4, although grouped as bandpass, has a rather high fraction rate for the low frequency bands compared to other band-pass clusters. This in line with our other observations that SWPC is quite biased toward low frequency connectivites.

Figure 7b shows the 8 clusters resulted from SWPC. Comparing Figure 7 with Figure 6, we can see that all the clusters resulting from SWPC are repeated in FBC results. This statement can be verified using Figure 7c. As seen in the aforementioned figure, all SWPC clusters have a high correlation with at least one of the FBC clusters. In contrast, 4 of FBC clusters are not visible in SWPC results, namely, clusters 4 through 8. All these clusters are either from band-pass or high-pass group and some show opposite patterns to each other pointing to an oscillating effect in connectivity patterns.

Looking at the comparison between TC and SZ fraction rate of FBC results (Figure 6 last row), we can see that, from the 8 comparisons, 5 are significant after correcting for multiple comparisons. TCs tend to stay more in clusters 1, 2 and 7, while SZs tend to stay more in clusters 5 and 8. As mentioned previously, clusters 1, 2 and 7 are related while cluster 5 is related to cluster 8. Lastly, if we look at the frequency profile of cluster pairs 5-8 (SZ>TC) in Figure 6 and compare them to cluster pair 2-7 (TC>SZ) we see that 5-8 clusters have relatively higher frequencies compared to the other pair. This is especially more visible for cluster 8 where the higher frequencies have higher fraction rate values.

## Discussion

In this preliminary work, we developed a new method (FBC) to estimate connectivity dynamics that are not limited to low frequency connectivity, or even to a specific choice of frequency. We first designed a toy example and showed that FBC enables us to estimate high-frequency changes in functional connectivity while typical SWPC might miss these changes. We then used FBC to analyze the FBIRN dataset and found 8 distinct connectivity states each with their own unique frequency profile. We showed that FBC is able to estimate the states resulting from SWPC in addition to some other states that go undetected using SWPC because of their higher frequency profile. In addition, FBC enables us to explore the frequency profile of connectivity patterns in the whole frequency range. i.e. using FBC we are able to comment on the frequency profile of each state. Applying this approach to real data reveals results in SZ that are consistent with, but extend, previous work and adds to our understanding of functional brain differences in this disorder.

### Activation frequency vs. connectivity frequency

As we mentioned in the introduction section, many studies have tried to explore the frequency profile of connectivity using different methods. The majority of these methods perform frequency tiling in the activity domain and then calculate connectivity. Then they proceed to make indirect inferences about the connectivity frequency profile. This is problematic as the relationship between the activity domain and the connectivity domain is unclear at best and depends heavily on the specific estimation method used. Many methods estimate connectivity (i.e. transform the activity domain into the connectivity domain) using highly nonlinear systems and therefore the frequency information is distorted in this transformation. For example, looking at the SWPC formula (Eq.1) we see that *μ*_*x*_(*t*) is subtracted from *x*(*i*) and the resulting value is divided by *σ*_*x*_(*t*). This part in itself distorts the frequency profile of *x*. In addition, [*x*(*i*) − *μ*_*x*_(*t*)]/*σ*_*x*_(*t*) is multiplied by [*y*(*i*) − *μ*_*y*_(*t*)]/*σ*_*y*_(*t*) to calculate correlation (Eq. 1). This step will further distort the frequency information. Therefore, using frequency information within the activity domain to infer frequency related information specific to connectivity is not straightforward. Some studies overlook this detail when studying connectivity frequency profile. For example, Li et. al. used frequency tiling in the activity domain to conclude that “oxygen correlation is band limited” (Li, et al., 2015). A similar issue can be found with a more recent paper where the talk about dynamic functional connectivity at specific bands (Luo, et al., 2019). In another type of study, Yaesoubi et al, used the wavelet transform to decompose the activity time series into different time-frequency bands and then calculate coherence. Several observations about connectivity frequencies are then made (Yaesoubi, et al., 2017). In our view, these kinds of statements can be misleading as the frequency profile is not directly studied in the connectivity space and therefore it does not enable us to make claims about the connectivity frequency profile. Rather they show that connectivity is caused by signals (i.e. activity) from specific frequency bands.

We believe to make correct claims about the connectivity frequency profile it is important to implement frequency tiling directly in the connectivity space. In addition, we should differentiate between the frequency profile of activity (estimated from time series themselves) and the frequency profile of the connectivity. The relationship between these two is not clear, so using knowledge about time series frequencies to make inferences regarding the connectivity frequency profile is not as straightforward as some studies suggest (Leonardi and Van De Ville, 2015).

### FBC performs frequency tiling in the connectivity domain

In this work, we have proposed an approach called FBC to estimate dFNC which does not make any assumption about the frequency profile of connectivity. This is in contrast with SWPC, which applies a low-pass filter when calculating dFNC. Note that we have filtered the BOLD time series using a band-pass filter (between 0.01 and 0.15) based on the existing literature (Niazy, et al., 2011). But to the best of our knowledge, no previous work has made the distinction between activity and connectivity frequency response and our work is the first one to do so. Because of this, we decided not to assume any prior knowledge regarding the frequency profile of connectivity, which resulted in considering 10 bands covering the entire sampled spectrum. Additionally, we explore the spectrum of activity time series and connectivity time series before filtering, i.e. w(t) and brought the results in the supplementary materials (Figure S13). As can be seen in this figure, the activity spectrum is clearly bounded between 0.01 and 0.15 Hz while the w(t) spectrum doesn’t seem to be bounded between any two frequencies. This point should be further explored in future complimentary works.

Using k-means clustering to summarize FBC results, we found that in addition to states resulted from SWPC, we can estimate some other states exclusive to FBC (see Figure 6). In addition, because of the frequency tiling nature of our method we are able to discuss the frequency profile of the estimated connectivity patterns. In many dFNC studies, it is reported that one dFNC state tends to be quite similar to the static connectivity (Damaraju, et al., 2014; Faghiri, et al., 2018). In this study, we see that there are actually 3 states that have strong low frequency profiles (states 1, 2 and 3; see Figure 6). State 1 is quite similar to sFNC resulted from SZ individuals while the other state, state 2, is similar to sFNC resulted from TCs (see Figure 7c for correlation between different states). One possible conclusion from this observation is that FBC enables us to distinguish between states that show more static-like behavior (states that show low-pass frequency profile) and others that are more dynamic (states that show high or band-pass frequency profile). This feature of FBC can be utilized so that we have a full picture of FNC in all its frequencies. i.e, study both static and dynamic FNC patterns simultaneously. As mentioned in the last paragraph we did not limit our connectivity spectrum, but this can easily be done. To do so, one can design filters to cover the desired bands.

### sFNC repeats in higher frequencies with lower cognitive control connectivity

Similar to the states reported in our earlier work (Damaraju, et al., 2014), several of the states resulted from the SWPC approach (Figure 7b) are quite similar to each other (states that show similar patterns to overall sFNC). We can see two states similar to these in FBC results too (namely states 2 and 4). Unlike the SWPC results, here we can see that while state 2 shows a very strong low-pass frequency profile, state 4 shows a more broadband frequency distribution. Again because of the frequency tiling of FBC, we have some added information in regard to the frequency profile of connectivity patterns that we can investigate. Cluster 4 has lower CC connectivity in general. One possible observation we can make here is that cluster 2 (which is quite similar to TC sFNC and has a very low-frequency profile) occurs in higher frequencies in the form of cluster 4 with smaller CC connectivity values. We speculate that CC (especially intraconnectivity within CC components) is lower in higher frequencies. This point should be examined in more depth in future work.

### SZ connectivity patterns have higher frequencies compared to TC connectivity patterns

SZ subjects have significantly higher fraction rate for cluster pairs 7 and 8 while TC subjects have significantly higher fraction rate for cluster pairs 4 and 6 (Figure 6). Looking at the frequency profile of these cluster pairs we can see that, cluster pairs 7 and 8 are in the high-pass group while pairs 4 and 6 are in the band-pass group. This seems in line with some previous studies where it was reported that in activity space SZs have higher power at higher frequencies compared to TCs (Abigail G. Garrity, et al., 2007; Alonso-Solis, et al., 2017). Unlike these studies, we have found alterations in the frequency profile of connectivity instead of activity. This observation has been made possible because of the unique feature of our proposed approach that allows us to study the frequency properties of connectivity directly. We believe these points grant further investigation.

### Weak connection between somatomotor and visual/auditory networks in SZ

FBC identified two unique states, 7 and 8, which are not visible via the SWPC approach. SZ spend significantly more time in these two states compared to TC (P<0.01, fdr corrected). Our approach enables us to intuitively capture the frequency profile and connectivity patterns of each state simultaneously which provide information that is not available to the SWPC approach. If we compare these states with states 4 and 6 (higher fraction rate for TCs compared to SZs) we see a difference in the connection between SM and sensory areas (AUD and VIS functional domains). These connections are strong (regardless of their sign) in states 4 and 6 while they are weak in clusters 7 and 8. Several studies have reported the presence of connectivity between motor and sensory regions in typical controls (D’Ausilio, et al., 2009; Londei, et al., 2010). A reduction in connectivity between these regions in SZ has been reported in the past years (Berman, et al., 2016; Kaufmann, et al., 2015).

### Dynamic connectivity highlights oscillations between two opposite patterns

Another observation we can make based on the FBC results (Figure 6) is that there are pairs of states (undetected by SWPC) that show opposite connectivity patterns. In our case, states 4 and 7 show opposite connectivity patterns of states 6 and 8 respectively. One interesting insight about these is that they exhibit a more high-pass frequency profile compared to the other states. This is possibly the reason that these states are not estimated using SWPC. As mentioned in the methods section, we used forward-backward filtering therefore It is unlikely that these opposite patterns are due to an alteration in phase caused by the analysis. Another explanation can be that these states are spurious estimations caused by the high-frequency nature of the filter banks as discussed by (Leonardi and Van De Ville, 2015). We cannot rule out this possibility completely but because these states show the opposite patterns of some other low-pass states this is unlikely. To further investigate, we removed the low frequency information of SWPC with different high-pass filters (forward-backward filtering) and then performed k-means clustering. Figure 8 shows the resulting clusters using different high pass filters. When removing the low frequency information, we obtain these opposite clusters (e.g. C3, Figure 8, quite similar to FBC cluster 6). Another possible explanation for these opposite patterns is that, similar to how the toy examples were designed, the dynamic connectivity patterns oscillate between two opposite patterns. Unlike the toy examples, the negative patterns (states 2, 4 and 8) show a higher frequency profile compared to their more low-pass counterparts, but this could reflect asymmetric oscillatory behavior (e.g. more rapid return from the negative patterns). This is an interesting observation; however, future work should be designed to further explore the possibility that connectivity indeed oscillates between two opposite connectivity patterns. In addition, the biological/clinical meaning of this view should be explored further.

**Figure 8.**
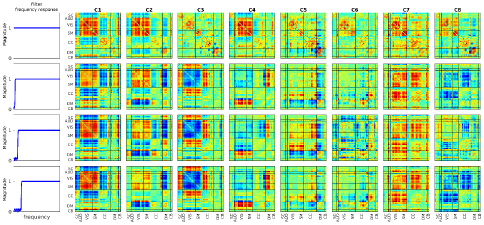
high-passed SWPC results. To verify FBC results we removed low frequency information of SWPC using different high-pass filters and then used k-means clustering. The first row is essentially unfiltered SWPC results while the next three rows are SWPC results using different filters. As seen in the last three rows we have clusters similar to FBC clusters in SWPC results too (clusters 3, 4 and 8). The only reason these clusters were not estimated using unfiltered SWPC is that SWPC frequency response function attenuates all frequencies above zero even in its band-pass.

### A systematic view of estimations

We believe that any analysis steps (including statistical estimators) can be viewed as a system and benefit from the extensive work done in the system design field (Bentley, et al., 2000). In this work, we examined SWPC and proposed an approximate system diagram for it (Figure 2.a). This way of thinking about SWPC facilitated our conclusion that SWPC has a low-pass filter inherent to its formula and led to our proposed method. The use of filter banks as proposed in the current work is only one possibility: for future works we can use custom h(t) functions to extract specific information. For example, one possible choice is the wavelet transform in the place of subsystem 2 (wavelet can be viewed as a system itself). Another possibility is to improve SWPC by replacing its h(t) with an IIR low-pass filter with very sharp transition and flat band-pass (in contrast with SWPC with a rectangular window where the frequency response of the filter is sinc like). This is in line with previous work which has suggested that SWPC window shape can be modulated to achieve better results (Mokhtari, et al., 2019).

### Robustness of the results in regard to different analysis choice

There are several choices we made which can impact the results. First, there is the choice of filter. We used Chebyshev type 2 filter for this analysis, but we also repeated all analyses with two other filter types (Butterworth and Elliptic) with matching characteristics. Figure S14 depicts the results for all 3 filter types. As can be seen in this figure, almost all the cluster repeats in all 3 filter types. The only difference is in cluster 7, where the fraction rate for Chebyshev filter is high-pass while the other two filters show a more band-pass fraction rate. This difference is probably caused by the difference in the frequency response of the filters. It is important to note that even though cluster 7 shows band-pass fraction rate for the last two filters, it has higher frequency profile compared to other band-pass clusters (clusters 4, 5 and 6). i.e. It started going down in higher frequencies.

Another parameter was window size used for estimating w(t). if Figure S4 through 12 we demonstrated that different window sizes with different cluster numbers show similar results. But to make sure the differences found between SZ and TC hold for different window sizes we di all the analysis with different window sizes (from 2TR to 60TR) and 8 clusters and performed all our statistical tests. As can be seen, almost all the results for different clusters hold, with the only distinction being cluster 7. This cluster is not estimated using higher window sizes, which leads us to believe that this cluster is showing very fast changes in variance and/or mean. Both Figures S14 and S15 show that we should be careful when talking about cluster 7 of our main results. On the other hand, Cluster 8 which shows an opposite pattern and form a pair with cluster 7 is repeated in both figures S14 and S15 with the same statistical test results (higher fraction rate in SZ). This observation reassures us that cluster 7 is a valid connectivity patter.

Finally, we repeated the analysis with different numbers of filters, while keeping the filter type, and cluster number the same as the main results of this paper. Figure S16, illustrate these results for different filter banks, from 5 to 12. Apart from clusters 4 and 8 (of the main results), all other 6 clusters are repeated in other rows with similar frequency profile. Cluster 4 is a little different in frequency profiles for filter number 5 and 8 compared to other filter numbers results. But the frequency profile, while categorized as low pas has a bump in middle frequency bands (unlike the first 3 clusters which are high only in very low frequencies). In addition, clusters in both columns 5 and 6 in Figure S16 are quite similar visually. Other different clusters are clusters in column 3 and 10. The reason behind this is possibly how the tiling has been done. When using filter numbers 5-7, we do not have good representation for the higher frequency (near normalized frequency 1) therefore the high-pass cluster is not estimated with this number of clusters. But for higher filter number this cluster has been estimated always (for filter number 8 through 12). Therefore, we think that this cluster is strong when we represent the highest frequencies more fairly.

### Limitations

The first limitation of the FBC approach is the subsystem used to transform activity space to connectivity space (Figure 2). In this system, we have used a window to calculate the mean and standard deviation. the problem arises when mean and std move faster than what the window can track. In this case, our estimations will be sub-optimal. To remedy this, we can use other instantaneous connectivity estimators suggested (Faghiri, et al., 2020; Shine, et al., 2015). Another limitation of this method is the possibility of noise contamination in the higher frequencies. This is certainly a valid issue, though our goal in this preliminary work was to refrain from making any strong assumptions about connectivity frequencies. The reason behind this decision was that we have no solid prior knowledge about connectivity frequency. In fact, to the best of our knowledge, this work is the first one that differentiates between activity and connectivity frequency profiles and previous works make assumptions about connectivity frequency using activity space information. Future investigators can utilize our proposed approach to study specific frequency bands more specifically. Another limiting feature is that in this study we have chosen to utilize our novel method to analyze a dataset that has been used extensively in many other studies. This allowed us to compare our results to the previously published work by using the same data and the same function for transforming from activity to connectivity (i.e. *w*(*t*)) as the previous work (Damaraju, et al., 2014). What we changed here is essentially the averaging step (Figure 2 subsystem 2) so that it does not remove frequency band information. We show that this provides a much richer source of information and additional insights into resting brain function in individuals with schizophrenia. One final limitation of this study is that, as shown in Figure 1, the estimated values at some bands might be non-significant. But we have used k-means to cluster the results which does not distinguish between significant and non-significant connections. To remedy this, in future work we can employ some methods that have a built-in statistical test. One such method can be change point-detection algorithms that have been increasingly applied to fMRI data (Jeong, et al., 2016; Xu and Lindquist, 2015). We can pair these methods with our estimation pipeline to find meaningful changes in the connectivity at different bands.

### Conclusion

In this work, we proposed a new approach to estimate dFNC called FBC. Our proposed approach does not make any strong assumption about connectivity frequency (unlike SWPC) and performs the frequency tiling in the connectivity domain. This is in contrast to previous works were frequency tiling was implemented in the activity domain. FBC aims to estimate connectivity in all frequencies and it enables us to investigate connectivity patterns frequency profiles. Using toy examples, we showed that FBC is able to even estimate high-frequency connectivities in addition to providing information about the estimated connectivity frequencies. Utilizing FBC, we analyzed an fMRI dataset including TC and SZ. Using FBC we found evidence of both static connectivity and time-varying states (typically identified with SWPC) in addition to some new connectivity states undetected by SWPC (possibly because of their high-pass nature). Finally, FBC points to a possible view of connectivity in which, data oscillate between two opposite connectivity patterns. This view should be further explored in future work.

### Data and code availability statement

Due to limitations imposed by the IRB we are unable to share the raw data, but it is possible to share the derived results. In addition, all the code used in this study will be included in the GIFT software and also shared upon direct request.

## Acknowledgments

Data collection was supported by the National Center for Research Resources at the National Institutes of Health (grant numbers: NIH 1 U24 RR021992, NIH 1 U24 RR025736-01, R01EB020407, P20GM103472, P30GM122734) and the National Science Foundation (1539067);

