## supplementary materials for "A unified approach for characterizing static/dynamic connectivity frequency profiles using filter banks"

**Choosing cluster number:**

To choose cluster number we evaluated a range of cluster numbers (2 through 25). Then we calculated the sum of within cluster distances for all cluster numbers and normalized all the values by the value calculated from using 25 clusters. Next we calculated how much each added cluster reduces the within cluster distance. The first cluster number that reduces this distance less than 0.05 was chosen as the cluster numbers. See Figure S3. Note that we also brought the results with different cluster numbers in Figures S4 through S12. All in all, we believe that cluster numbers should be treated more like fast Fourier transform length (especially in this approach). Results show there is no “best” cluster number and increasing and decreasing this number changes the resolution of the effect we can explore. Proving this more comprehensively is beyond the scope of this manuscript and will be explored in a future study.

**The effect of window size:**

To explore the effect of window size, we used different window sizes for calculating FBC and then performed the clustering with different cluster numbers (Figures S2 through 10).

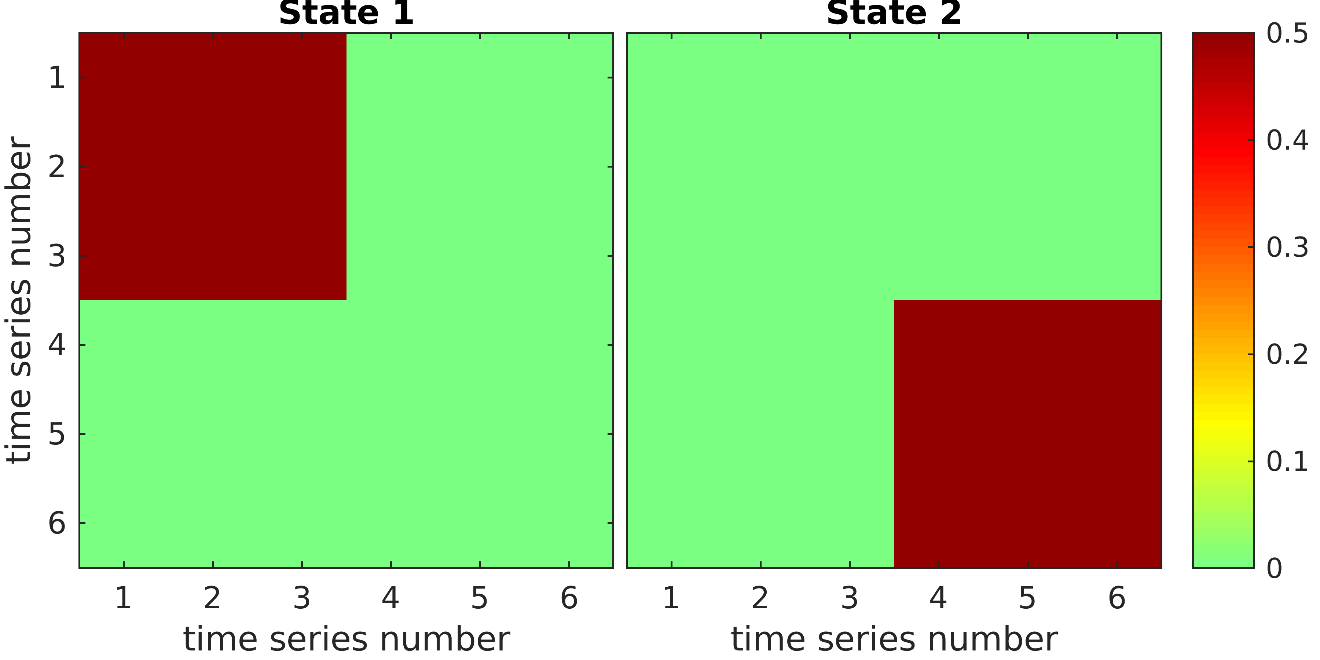

Figure S1 Correlation amplitude matrix ($\boldsymbol{A(t)}$) for the two states

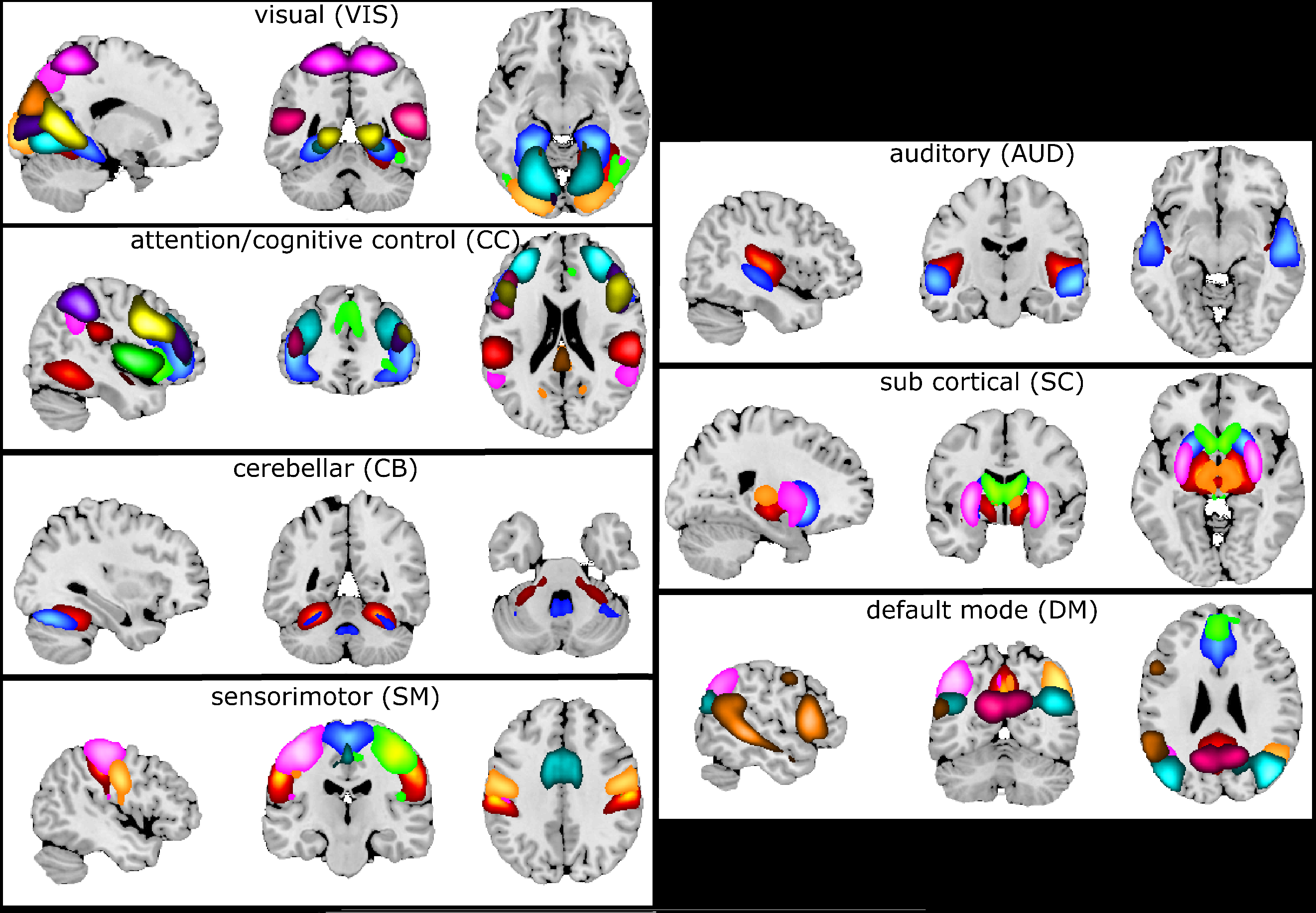

Figure S2 Spatial maps for different functional domains. Group ICA was used to calculate 100 maximally spatial independent components. These 100 component spatial maps were visually inspected and 48 of them were chosen and grouped into 7 functional domains.

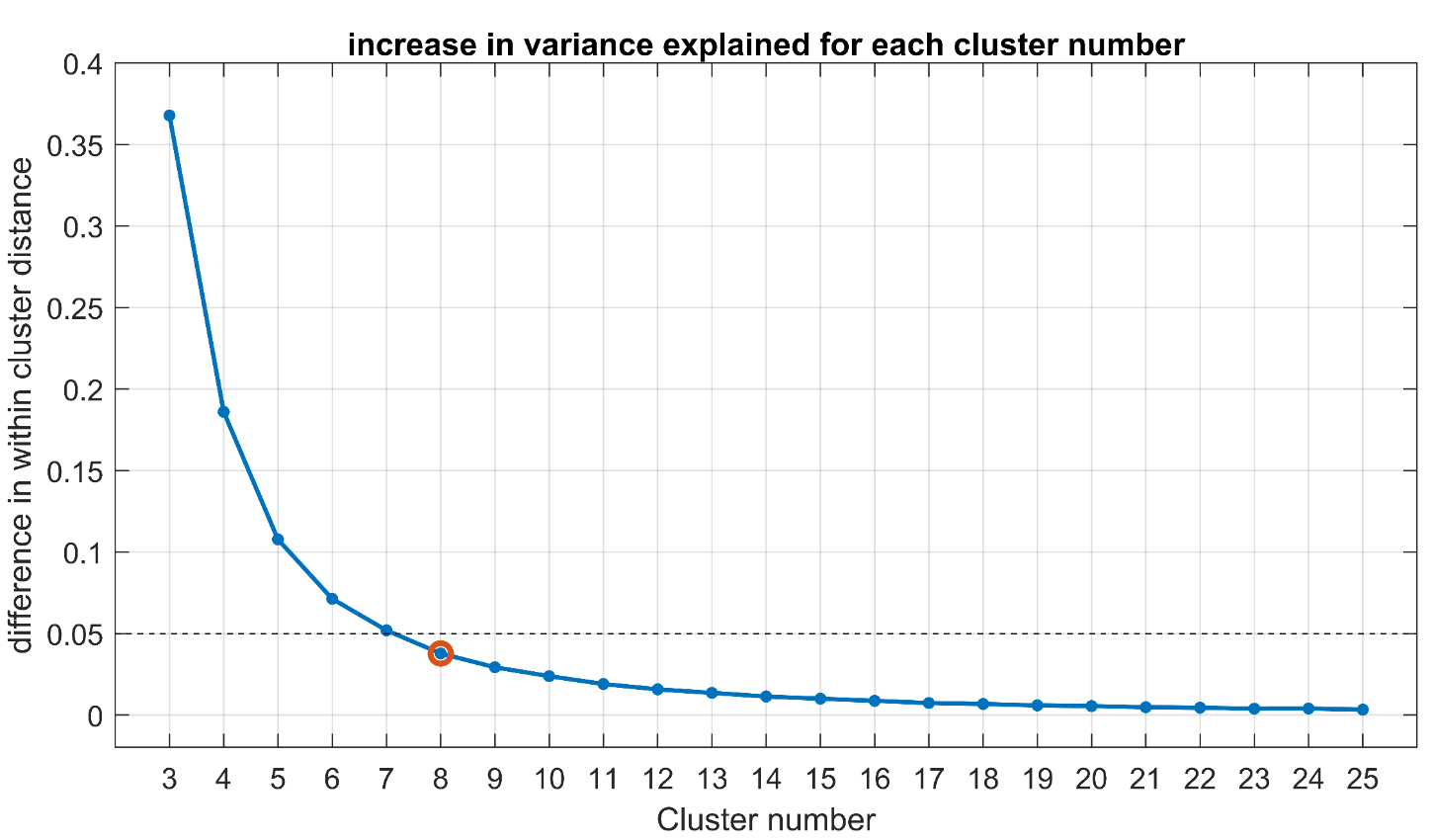

Figure S3 decrease in within cluster distance with each added cluster. As can be seen here, the first cluster with a difference below 0.05 (normalized value) is cluster 8. Therefore, we selected 8 as the cluster number for further analysis.

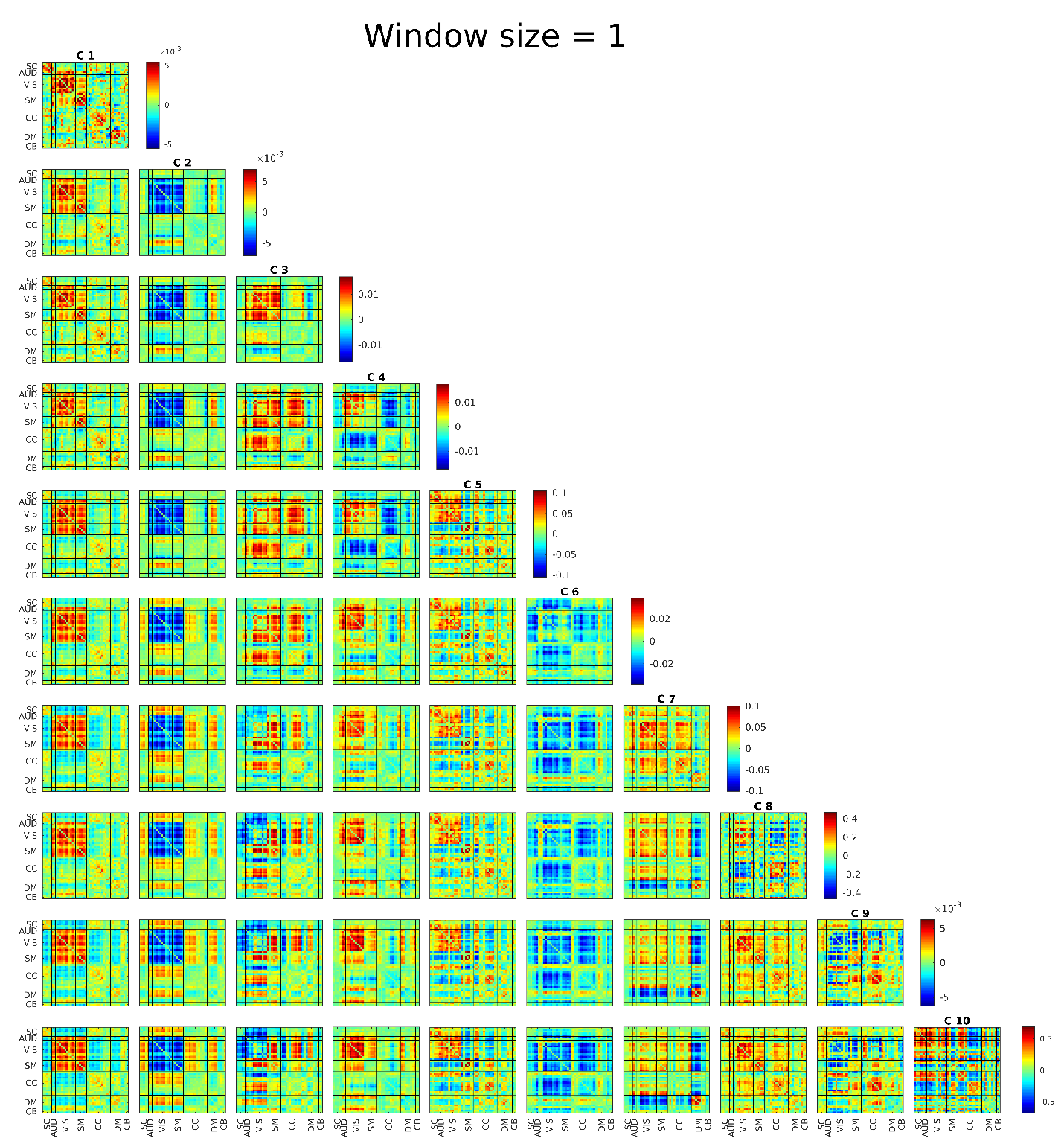

Figure S4. FBC results for window size equal to 1TR (2sec)

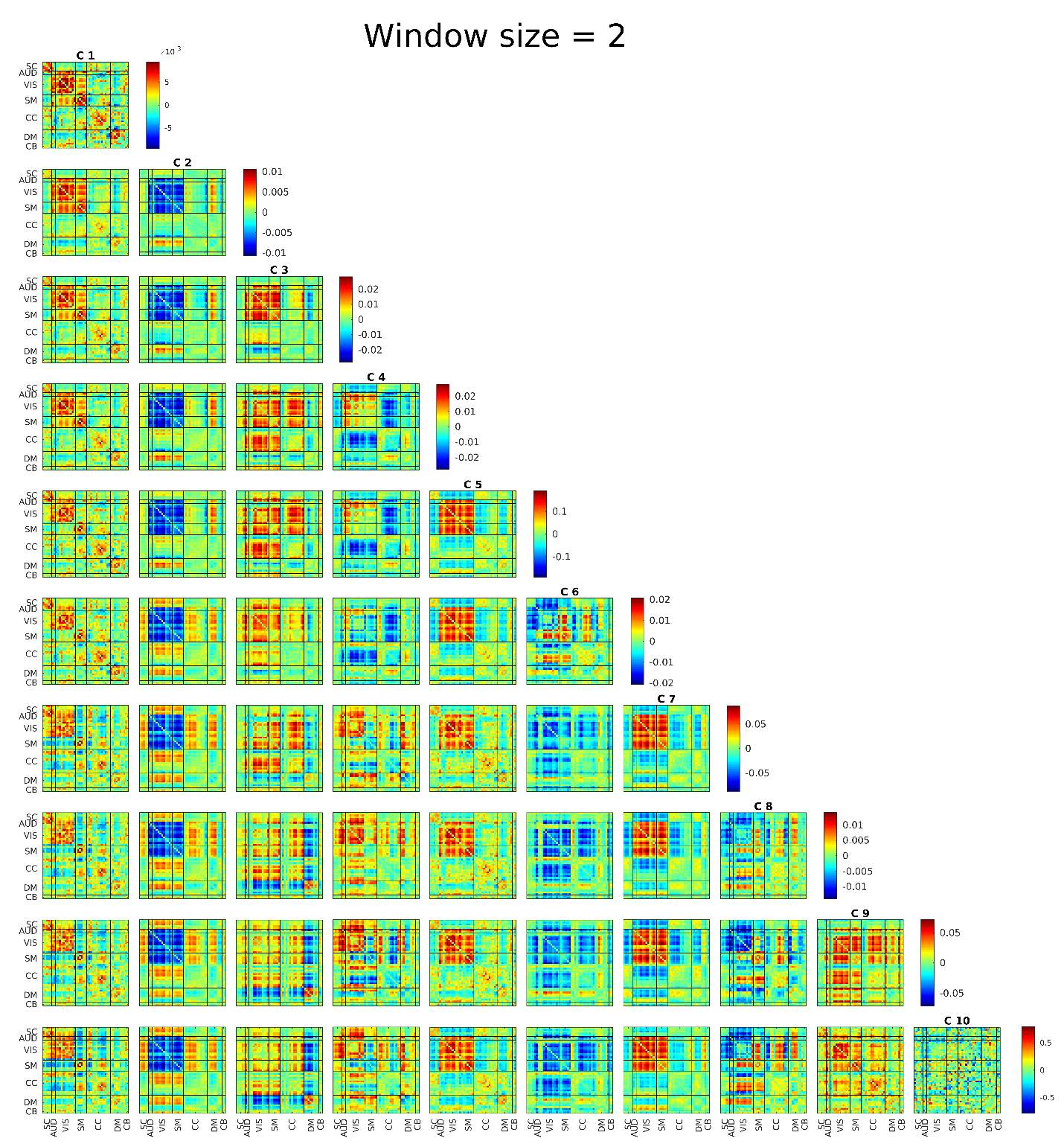

Figure S5 FBC results for window size equal to 2TR (4sec)

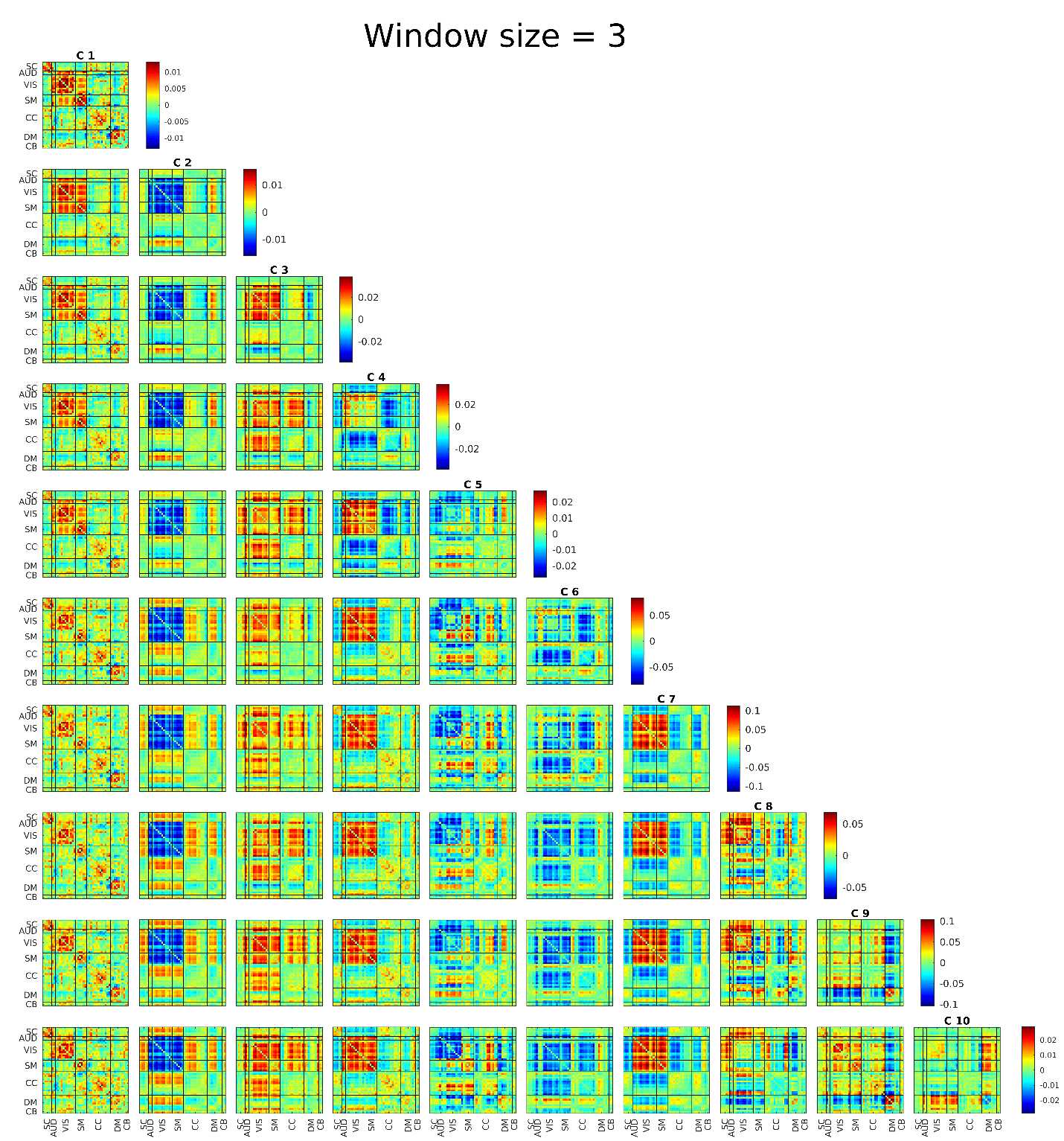

Figure S6 FBC results for window size equal to 3TR (6sec)

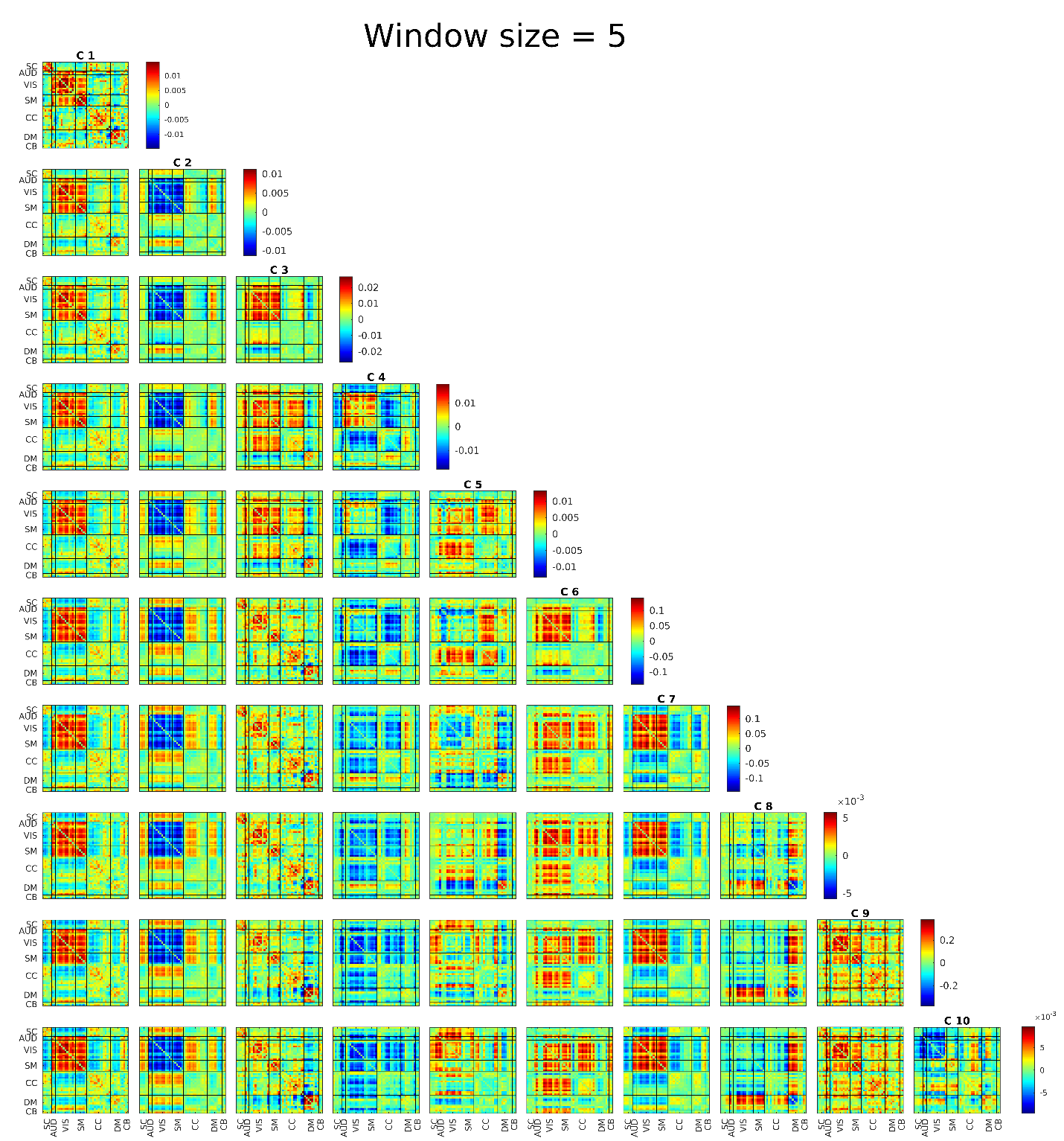

Figure S7 FBC results for window size equal to 5TR (10sec)

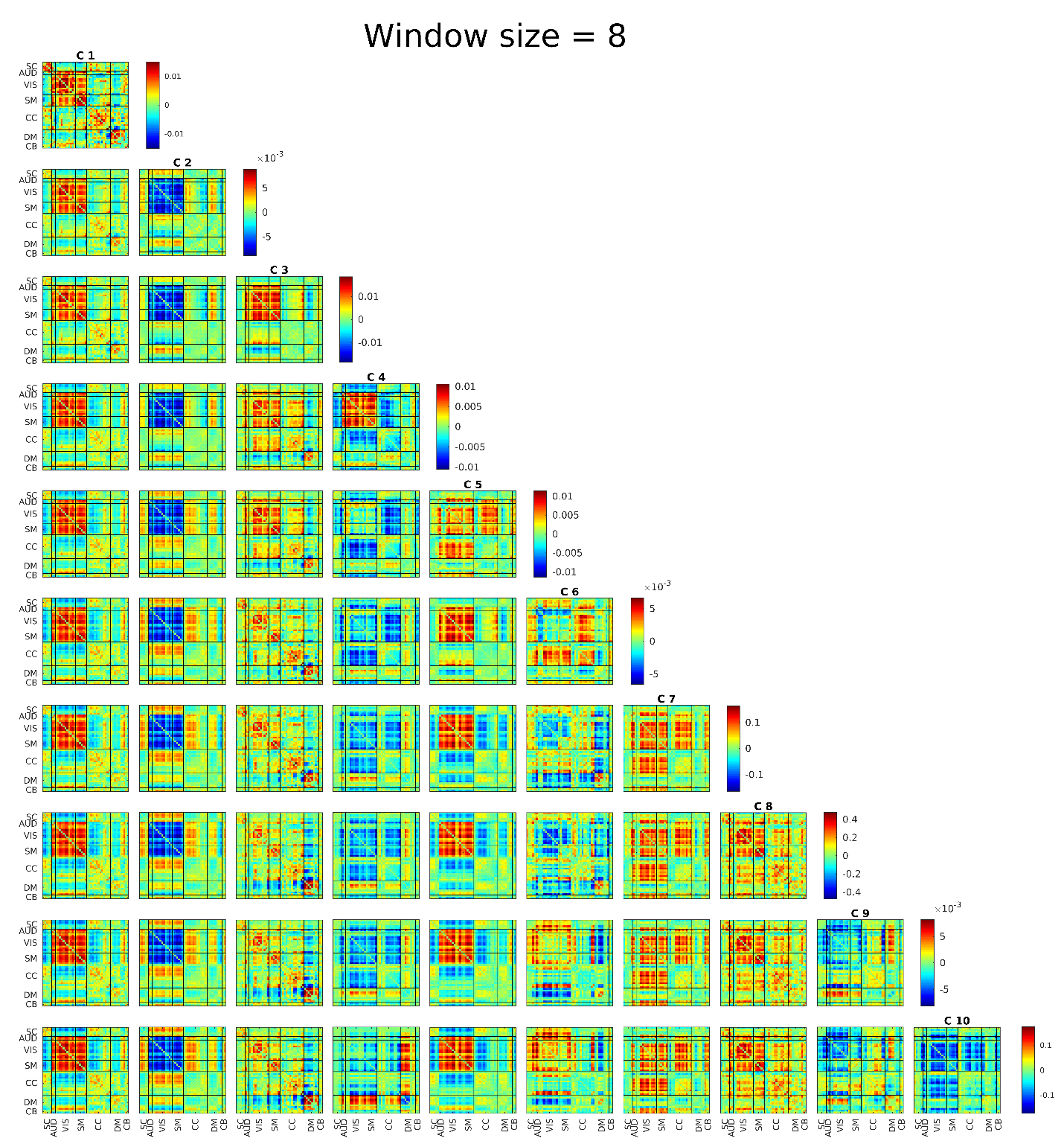

Figure S8 FBC results for window size equal to 8TR (16sec)

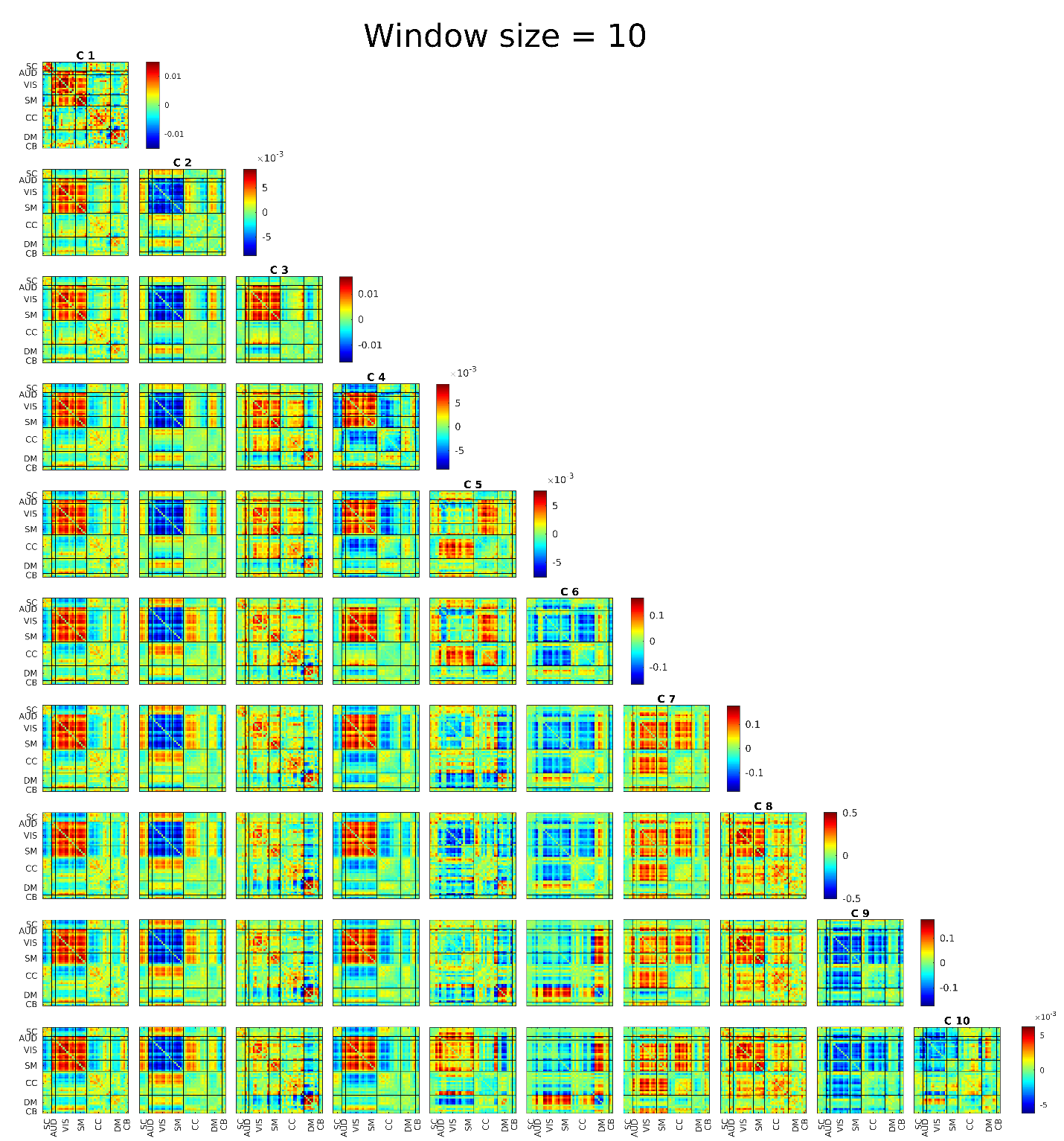

Figure S9 FBC results for window size equal to 10TR (20sec)

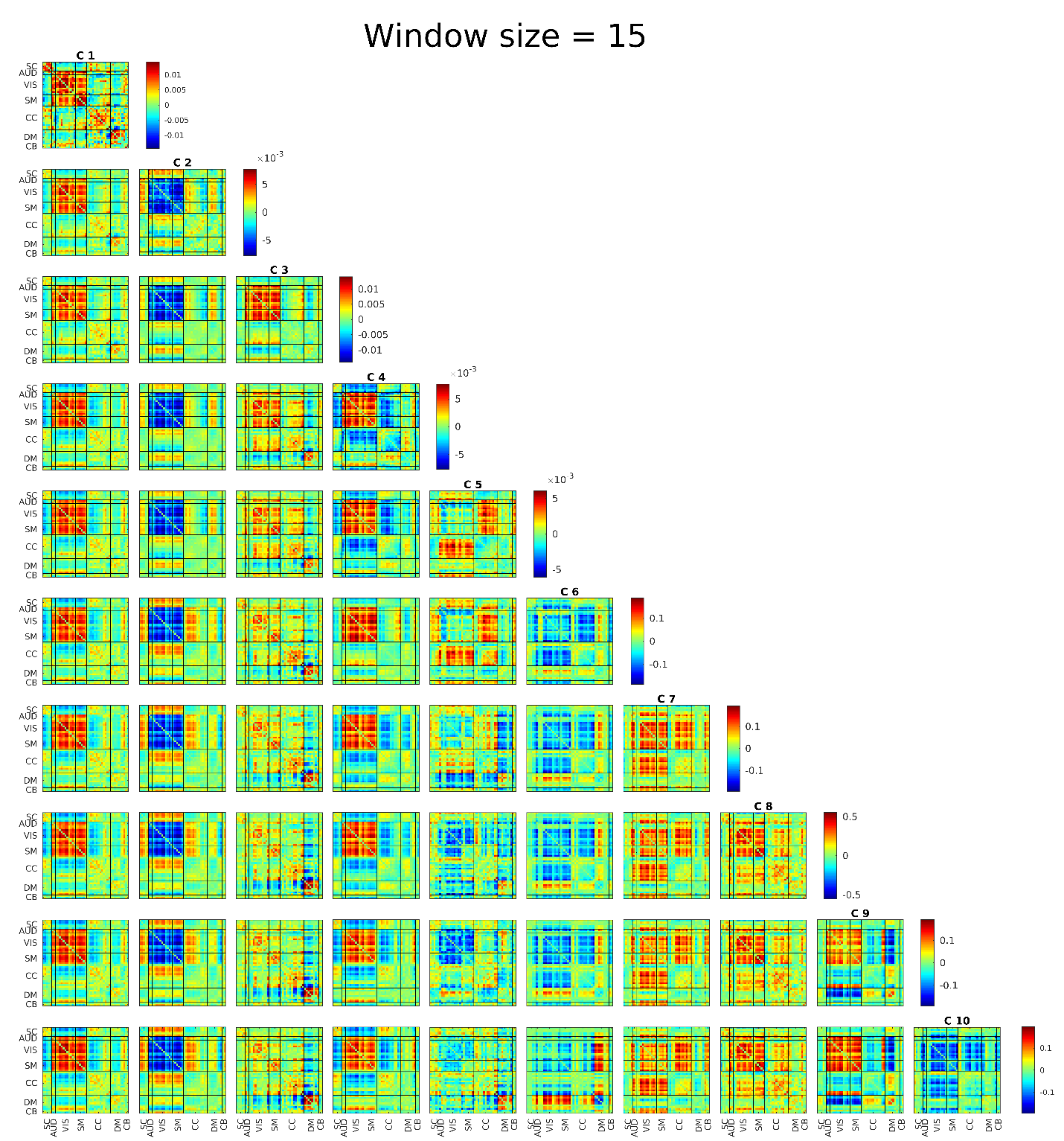

Figure S10 FBC results for window size equal to 15TR (30sec)

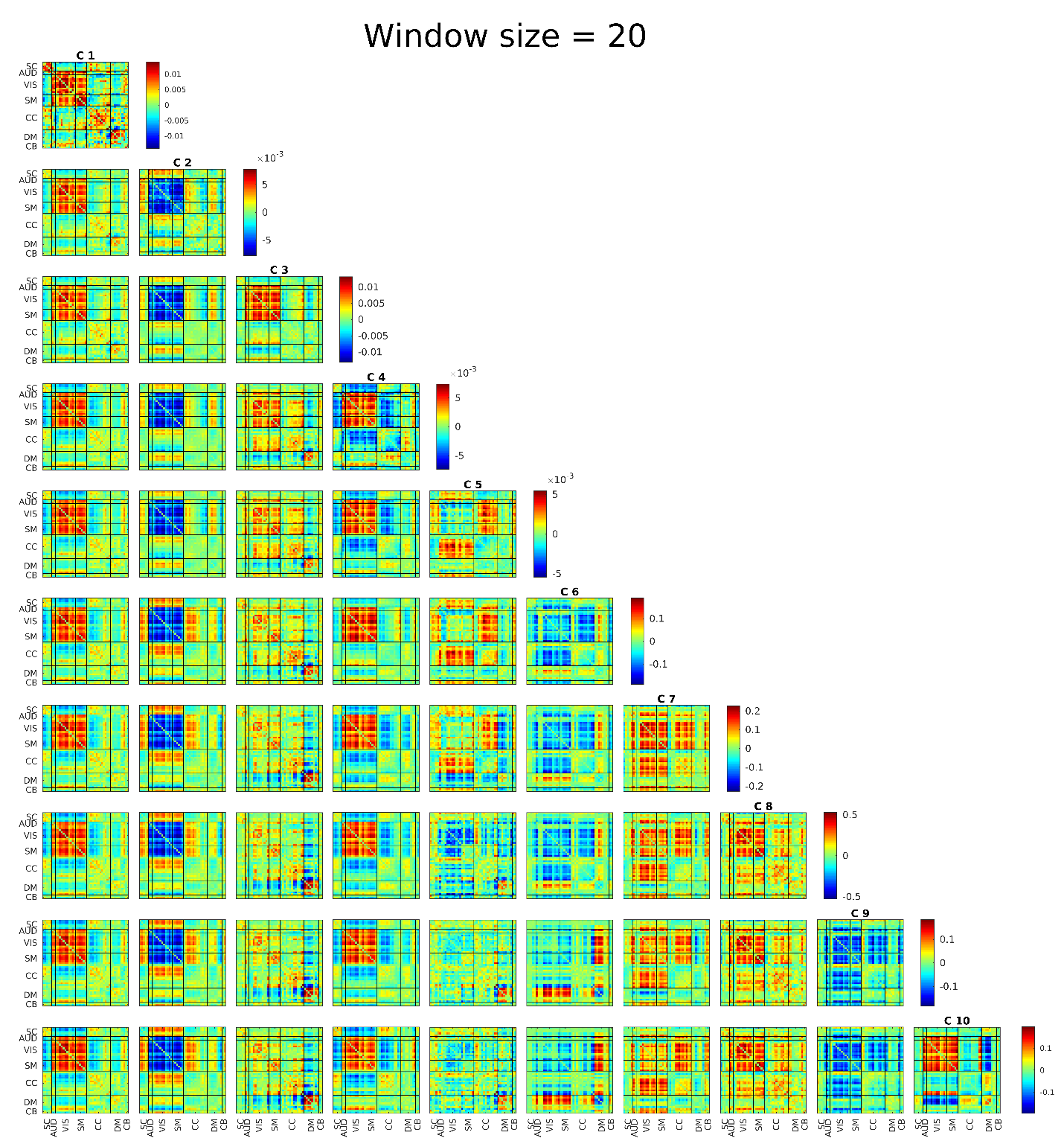

Figure S11 FBC results for window size equal to 20TR (40sec)

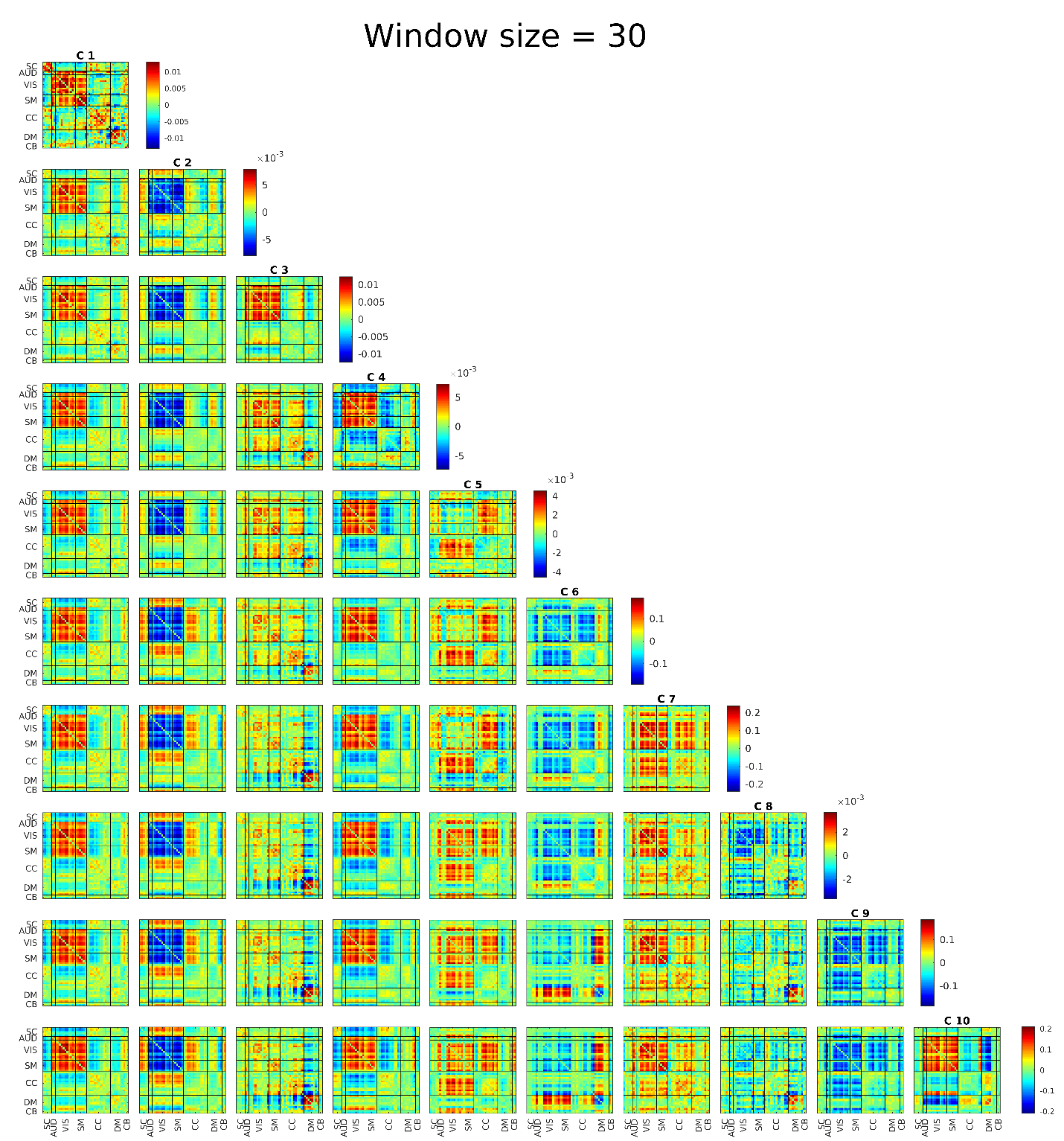

Figure S12 FBC results for window size equal to 30TR (60sec)

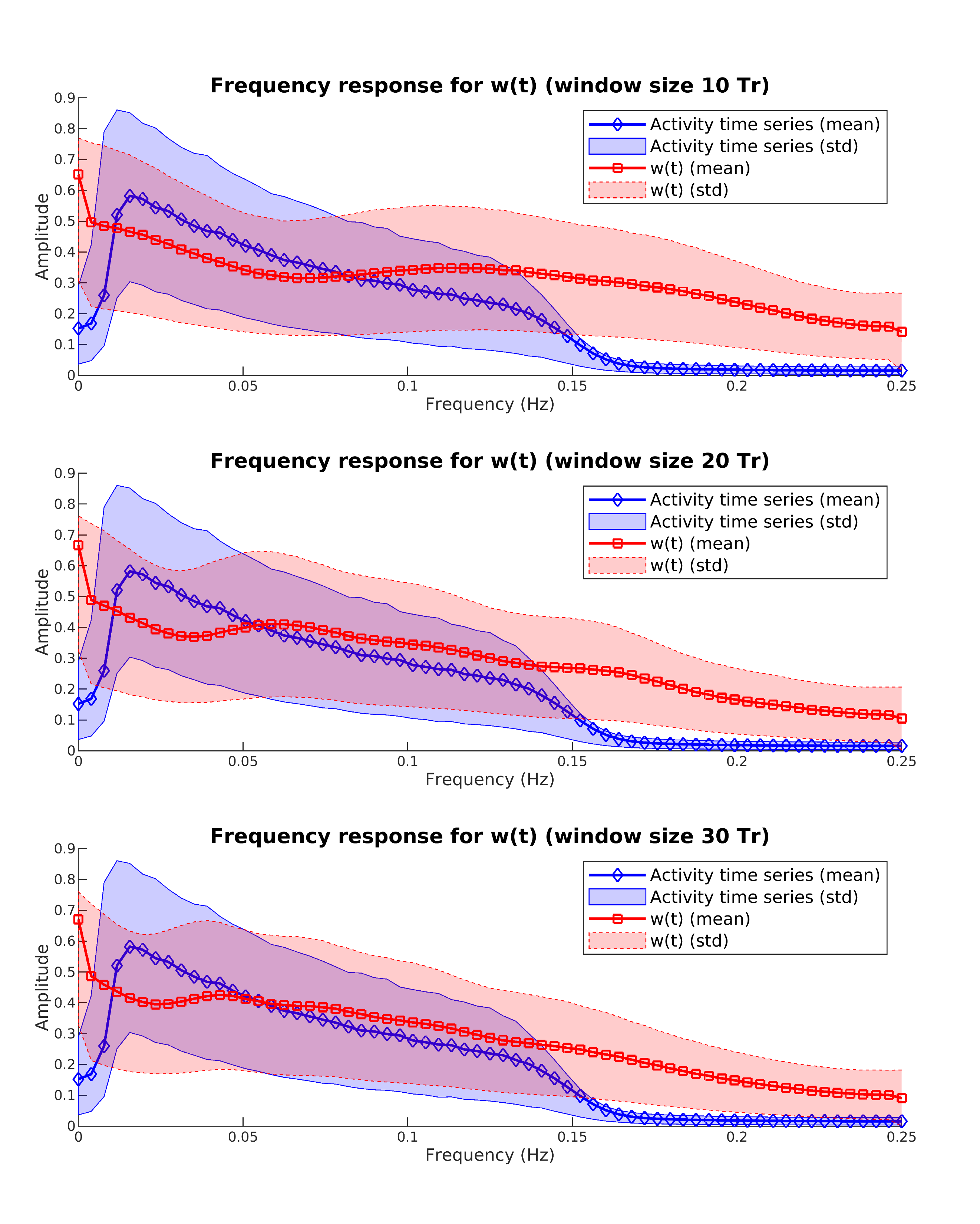

Figure S13. Average frequency response of activity time series and w(t), i.e. connectivity, time series. AS can be seen here, the activity time series is bounded between 0.01 and 0.15. This was to be expected as we filtered the activity to be in this range. But the connectivity time series do not seem to be bounded and it goes down monotonically.

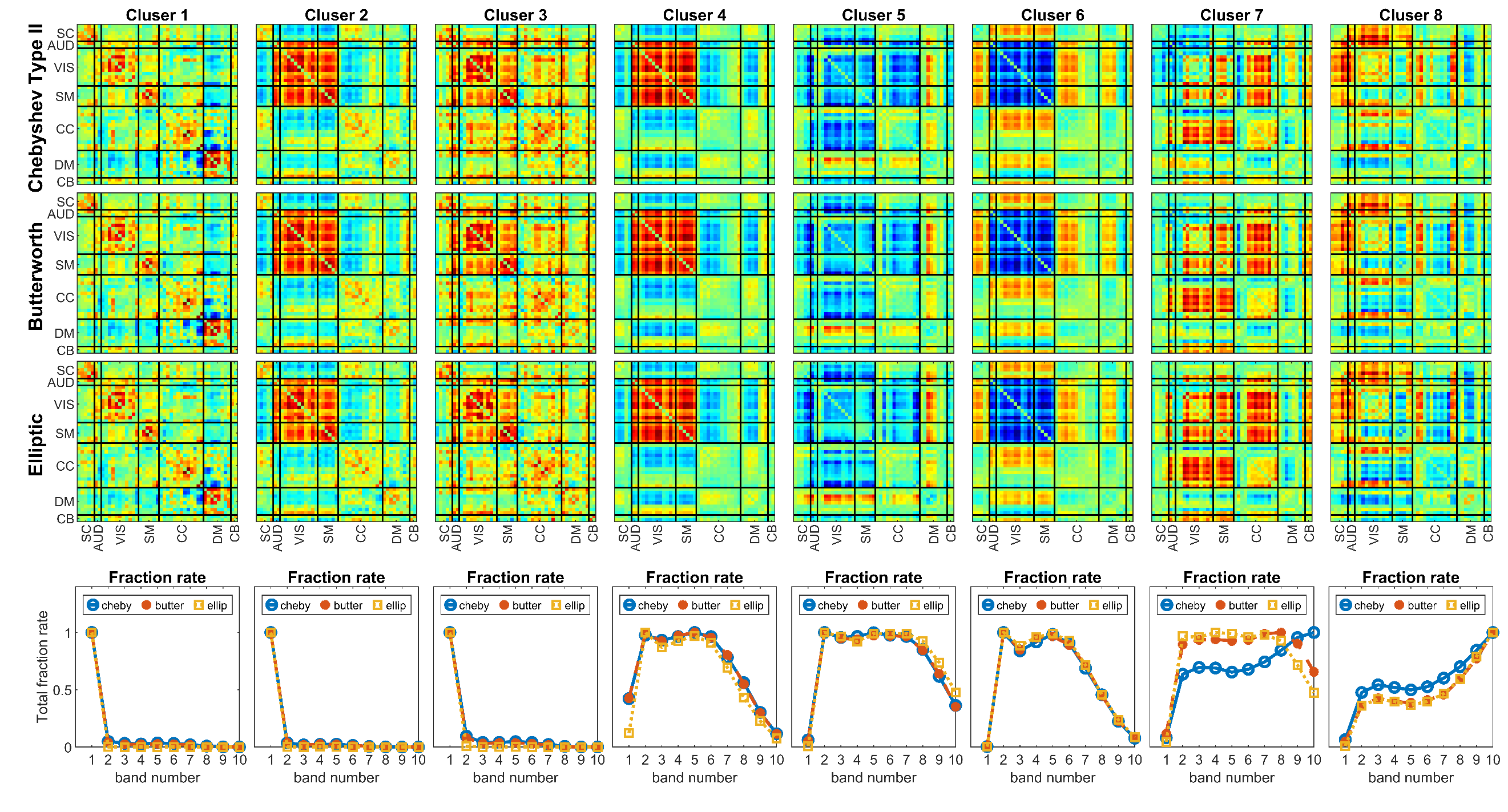

Figure S14. Results using 3 different filter types with similar characteristics. The first row shows essentially the main results of the paper while the second and third row, demonstrate the results using Butterworth and elliptic filters. The fourth row shows the frequency profile (i.e. fraction rate for each band) of each cluster with each filter. All the clusters are quite similar in both their centroids patterns and their frequency profile. The only exception is cluster 7 where the frequency profile is a little different. The difference is not much and we think it is because of the difference in how sharp each filter transit from pass-band to stop-band for these filters.

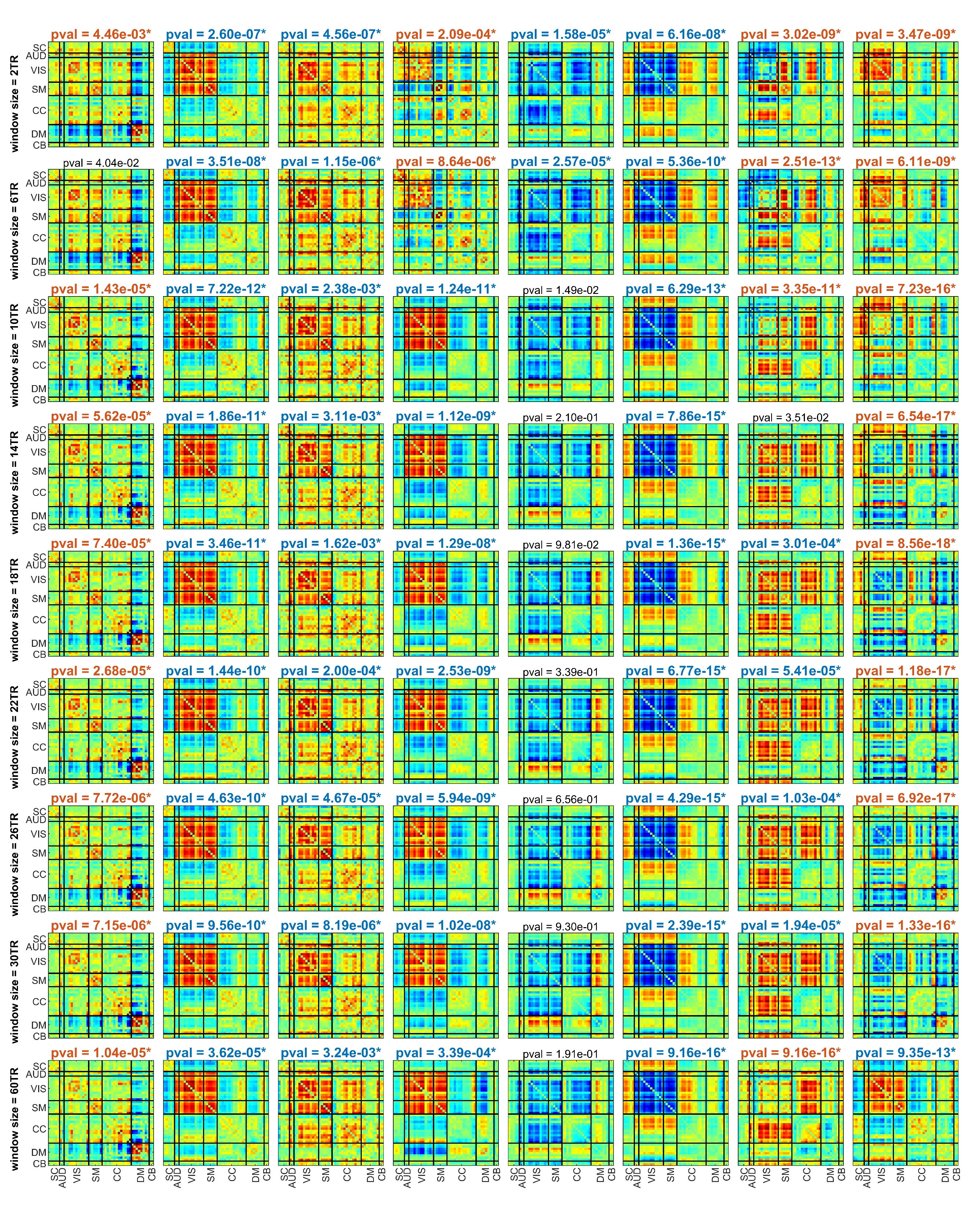

Figure S15. Statistical difference between SZ and TC across different window sizes. Blue title means TCs have significantly higher fraction rates while the red fonts mean SZs have significantly higher fraction rates (fdr corrected, p<0.01). non-bold black fonts mean non-significant.

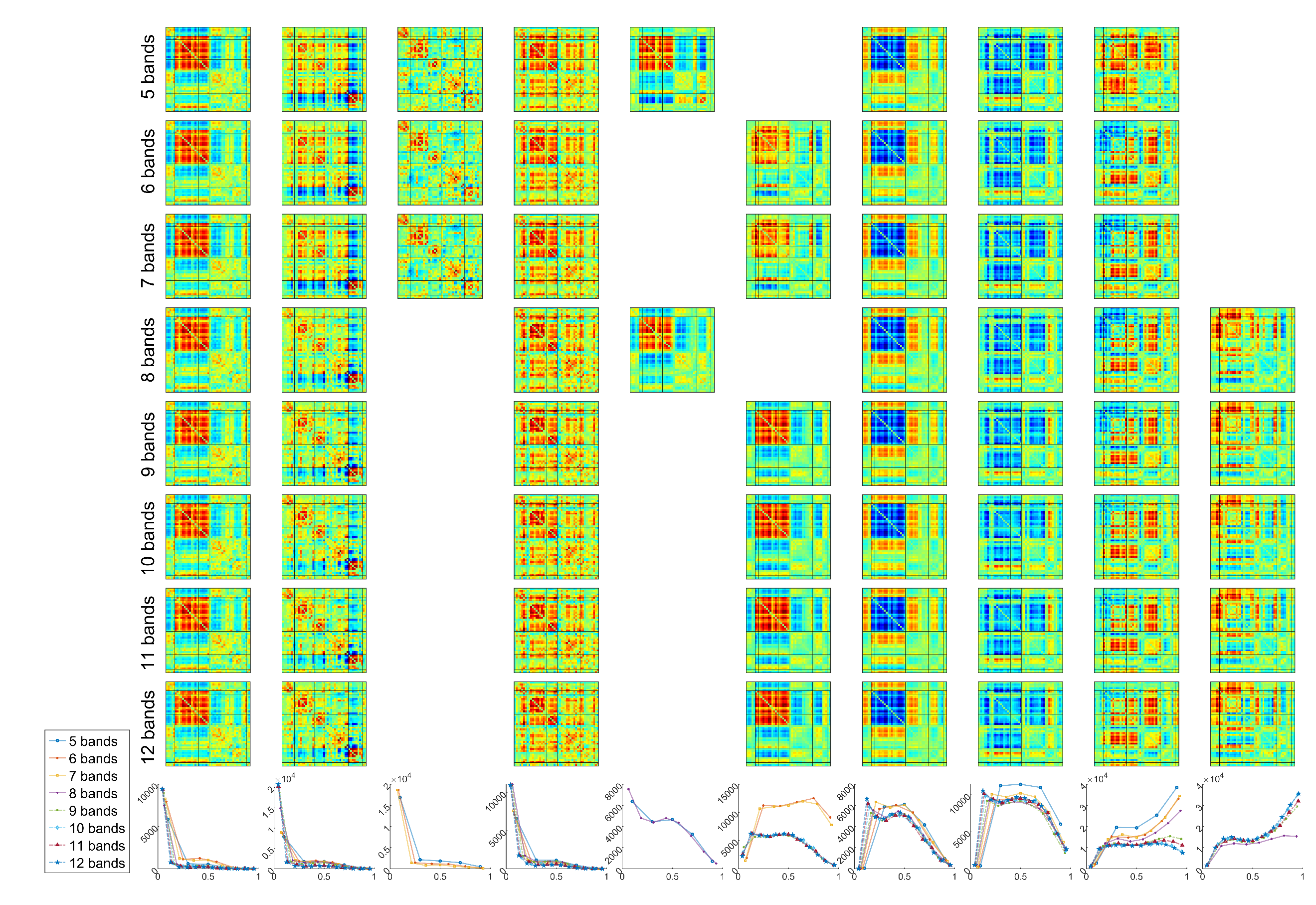

Figure S16. Clustering results using different filter numbers. Rows 1 to 8 show the results from 5 filters to 12 filters respectively, while the last row shows the frequency profiles of each cluster. The clusters are matched in each column and put in a separate column if no match was found for them. 6 clusters of the main results (filter number 10) repeat in all filter number with very similar frequency profiles (columns 1, 2, 4, 7, 8, and 9 in this figure). The only discrepancy is between is in columns 3-10 and columns 5-6. Columns 5-6 show very similar centroids patterns and while their frequency profile is somewhat different, column 5 shows a bump in middle frequency ranges. This is unlike other low pass clusters where the maximum is in the first band and fraction rate goes down for higher frequencies monotonically. The two other differences are probably caused by the difference in how the tiling is done between different filter numbers. If we have higher number of filters, different band ranges will have more direct and specific representations. For example, we can say that for filter numbers above 8 high frequencies are represented more therefore we are seeing column 10 clusters with these filter numbers. This statement is validated by the fact that the frequency profiles have higher values for high-frequency bands for higher number of filters. Compare purple with dot marker (8 bands) to blue dashed line (12 bands) in the last row.
